# Temporal control by co-factors prevents kinetic trapping in retroviral Gag lattice assembly

**DOI:** 10.1101/2023.02.08.527704

**Authors:** Yian Qian, Daniel Evans, Bhavya Mishra, Yiben Fu, Zixiu Hugh Liu, Sikao Guo, Margaret E. Johnson

**Author notes:** these authors contributed equally (Yian Qian, Daniel Evans).

## Abstract

For retroviruses like HIV to proliferate, they must form virions shaped by the self-assembly of Gag polyproteins into a rigid lattice. This immature Gag lattice has been structurally characterized and reconstituted *in vitro*, revealing the sensitivity of lattice assembly to multiple co-factors. Due to this sensitivity, the energetic criterion for forming stable lattices is unknown, as are their corresponding rates. Here, we use a reaction-diffusion model designed from the cryo-ET structure of the immature Gag lattice to map a phase diagram of assembly outcomes controlled by experimentally constrained rates and free energies, over experimentally relevant timescales. We find that productive assembly of complete lattices in bulk solution is extraordinarily difficult due to the large size of this ∼3700 monomer complex. Multiple Gag lattices nucleate before growth can complete, resulting in loss of free monomers and frequent kinetic trapping. We therefore derive a time-dependent protocol to titrate or ‘activate’ the Gag monomers slowly within the solution volume, mimicking the biological roles of co-factors. This general strategy works remarkably well, yielding productive growth of self-assembled lattices for multiple interaction strengths and binding rates. By comparing to the *in vitro* assembly kinetics, we can estimate bounds on rates of Gag binding to Gag and the cellular co-factor IP6. Our results show that Gag binding to IP6 can provide the additional time-delay necessary to support smooth growth of the immature lattice with relatively fast assembly kinetics, mostly avoiding kinetic traps. Our work provides a foundation for predicting and disrupting formation of the immature Gag lattice via targeting specific protein-protein binding interactions.

## Introduction

All retroviruses, including HIV, must form new virions that can bud out of the plasma membrane of the infected host cell (1). These virions are comprised of the retroviral polyprotein Gag, genomic RNA (gRNA), and additional co-factors(1). Gag is the primary structural component of the virions, initially assembling an immature lattice that is bound to the membrane and forms a tri-hexagonal organization as revealed by cryo-electron tomography (cryoET)(2, 3). Once the virion has budded, the Gag polyprotein is cleaved by its attached proteases(4) and reassembles into the mature viral capsid (5). The stability of the immature Gag lattice appears tuned to ensure successful assembly while also ensuring the remodeling necessary to transform from the immature lattice to the infectious mature capsid(6, 7). Indeed, maturation inhibitors are a promising strategy for antiviral drugs that function by over-stabilizing the immature lattice(7-9). Assembly of the immature lattice can be reconstituted *in vitro*(10, 11), but because the immature lattice does not assemble from the Gag monomers alone(10), it is not known how the energetics and cooperativity of Gag-Gag contacts, Gag contacts with co-factors, or their timescales of binding drive stable lattice formation. Here, we develop a coarse-grained model of Gag assembly to quantify assembly pathways as a function of Gag-Gag binding rates and stability, ultimately designing time-dependent protocols that allow us to robustly assemble immature lattices with varying kinetics. These models on their own and with comparison to *in vitro* kinetics(11) predict how time-dependent control of co-factor binding can induce (or disrupt) stable assembly of the immature Gag lattice.

A primary challenge in understanding and predicting the assembly of the immature Gag lattice is its dependence and sensitivity to a range of co-factors. The minimal conditions to assemble the virus-like lattice in solution *in vitro* requires Gag along with at least one negatively charged molecule like RNA or the cellular small molecule inositol hexakisphosphate (IP6) (12-14). Gag does form stable dimers on its own, which are further strengthened with co-factors present (15). Thus, it is the formation of the higher order contacts in the immature lattice that must be distinctly ‘switched on’ by interactions with co-factors, at least in part by inducing conformational changes into the Gag proteins(10). Combining RNA and IP6 together works synergistically to promote and accelerate immature Gag lattice assembly, indicating they have somewhat complementary roles in stabilizing the lattice (11). With just RNA or just IP6, assembly takes ∼1-2 hours at 50*μ*M Gag, whereas with both, it can proceed in seconds (11). Indeed, IP6 promotes formation of infectious Gag virions *in vivo* by stabilizing the immature lattice (6). IP6 coordinates with the hexamer contacts in the Gag lattice in a 1:6 stoichiometry, stabilizing Gag into an assembly-competent form for the immature Gag lattice (15). The immature Gag lattice also contains another set of interfaces that form a trimeric cycle that can provide additional stability when brought together via the dimer and hexamer assembly, although it is likely that these smaller interfaces are relatively weak (16). Through our computational models, we can reach the seconds to minutes timescales of co-factor mediated assembly, where robust growth and a high yield of virions forms within minutes if a high enough concentration of RNA and especially IP6 are present (11). We can thus directly test how the relative strengths and speeds of the different Gag-Gag interactions, as they would be triggered in time by co-factors, controls lattice assembly.

Another challenge with the immature Gag lattice is its size; a completed spherical lattice contains ∼4000 monomers, which is an order of magnitude higher than most computational models of self-assembly. There are two major issues that arise with this larger size. The first is simply that the models become more computationally expensive to characterize across varying parameters compared to completed assemblies that contain 12 (17, 18), 60 or even 100s of subunits (19, 20). For smaller systems, systems of rate equations can be constructed to efficiently characterize phase diagrams and assembly pathways as thermodynamic and kinetic parameters are varied (18). Spatial simulations using Brownian dynamics simulations can also be relatively efficient for characterizing assembly pathways in systems with ∼100s of subunits (19, 21), but 1000s of subunits are rare (22). Non-spatial simulations of large HIV-like assemblies must limit the types of assembly pathways followed(23). The second major issue, however, is that with more subunits required to complete the assembly, the rate of nucleating new lattice structures must be correspondingly slowed to allow elongation of existing nuclei to completion (24). Otherwise, assemblies become kinetically trapped in intermediates that cannot readily combine to form larger structures, as monomers are fully depleted, or starved (25). The parameter regimes that can support assembly will be significantly compressed relative to assemblies with fewer subunits. Hence, the Gag lattice is intrinsically primed to be prone to kinetic traps due to its size.

Kinetic trapping, which can dramatically slow the formation of productive and complete equilibrium assemblies, sometimes beyond experimentally relevant timescales, is a consequence of the nonequilibrium nature of the self-assembly process. Subunits are initially unbound, and the pathways they follow towards reaching the more stable equilibrium structures determine whether incomplete intermediates consume the unbound monomers. For such a large lattice, how does HIV avoid kinetic traps? *In vivo*, we identify two main strategies that can will help avoid kinetic traps and productively assemble virions. First, subunits do not appear in bulk in the cytoplasm but are produced at a rate that slowly increases their concentration in time and space (26). Second, as detailed above, co-factors can ‘activate’ the Gag monomers such that lattices only nucleate and grow when they have bound RNA. Both these mechanisms introduce additional timescales that can reduce nucleation while still ensuring growth of viruses. Theory and modeling have shown how, although kinetic traps for smaller assemblies can be reduced by optimizing energetic parameters to moderately weak strengths (19, 25), similar strategies to those used by the cell that introduce new timescales can significantly improve yield. Cooperativity during growth that disfavors formation of new nuclei by modulating binding sites helps avoid traps (27), as can variable addition of monomers (28), and allosteric activation (29). *In vitro* studies of HIV-1 immature lattice assembly in solution showed how the stoichiometry of the co-factors relative to the Gag monomers tune the kinetics of lattice assembly, indicating that co-factor binding can act as a rate-limiting timescale in assembly (11). In our work here, we therefore use the theory of diffusion-influenced reactions (30-32) to derive a timescale that could effectively represent these same biologically relevant processes used to suppress trapping, by directly controlling the concentrations of monomers with time. Our simulation results illustrate how even this single additional timescale can effectively eliminate the trapped intermediates to promote successful assembly.

A key advantage of the structure-resolved reaction-diffusion modeling approach we use here is that the timescales and assembly yield are controlled by biochemically measurable rates and free energies, and thus the predicted timescales we define compare directly to experiments. Recently, coarse-grained models of the immature HIV lattice have shown how IP6 stabilizes lattice assembly and speeds up binding with more molecular details, in ways that may promote defects in the structures (33). These coarse-grained models built on earlier work characterizing the specificity of Gag contacts needed for proper assembly (34), and the role of RNA and membrane binding in stabilizing Gag assembly (35). These models successfully captured morphological variations in assembled structures and explicit co-factor interactions but are not able to map the kinetics of binding and assembly directly to experimental timescales, and the strength of protein-protein interactions can be similarly difficult to map to experimental K_D_ values. Our model is thus complementary, producing assembled structures that are similarly derived from experiment and build from the monomeric building blocks, while providing direct observations of tunable assembly kinetics over longer (minutes) timescales. We are thus able to estimate rates for Gag binding to the co-factor IP6 by comparison with the *in vitro* kinetic data (11), and set bounds on the rates of Gag-Gag binding interactions.

In this paper, we first describe the construction of our computational model of Gag self-assembly from cryoET structures and validate the kinetics and equilibrium of our simulations with stepwise comparisons to theory. We tested a range of binding free energies and rate constants for the distinct Gag-Gag binding interactions of dimer, hexamer, and trimer contacts. We identified regimes where assembly could be eliminated, where a two-phase equilibrium of dilute monomers and partial assemblies formed, and where kinetic trapping onset. Kinetic trapping was predominant, with our structural analysis quantifying the frequent irregularity of the ensuing lattice growth. To mimic the time-dependent activation of Gag by co-factors, we used a titration rate to slowly increase the Gag concentration, finding that when the rate was slow enough, we produced robust and regular lattice growth. We therefore derived a titration rate dependent on the underlying Gag assembly kinetics, showing that it is remarkably successful in promoting productive lattice assembly. Finally, we connected our model to *in vitro* light scattering kinetics of Gag assembly driven by co-factors RNA and IP6. We were able to estimate bounds on binding rates between Gag and IP6, and Gag-Gag using relatively simple theoretical arguments. We conclude by discussing the future extensions and applications of this integrative modeling approach.

## Methods

### Model construction

Our model consists of coarse-grained representations of Gag protein monomers derived from the cryoET structure (16) of the immature Gag lattice (Fig 1a). Each Gag is a rigid body with a center of mass (COM) point and five distinct binding sites (Fig 1B). The center-to-center distance between two hexamer substructures is 7.7 nm, the final sphere radius R=50nm, and the complete lattice contains ∼ 3700 Gag proteins. There are 3 types of binding interactions: the homo-dimer interaction, the hexamer interaction mediated by the two distinct hexamer sites, and the trimer interaction mediated by the two distinct trimer sites. When two molecules bind via these specific interaction sites, they adopt a pre-defined orientation that ensures assembly into the experimental cryoET structure. To derive our model of Gag-Gag interactions from the cryoET structure, we must correct for the small variations of the 18 monomer positions within the lattice. Without corrections, our monomers would assemble spherical fragments with distinct curvatures and thus significant defects due to slight changes of orientations between hexamer, trimer, and dimer contacts. Thus, we symmetrized the Gag lattice to ensure a single curvature across all contacts. See SI for mathematical details.

**Figure 1.**
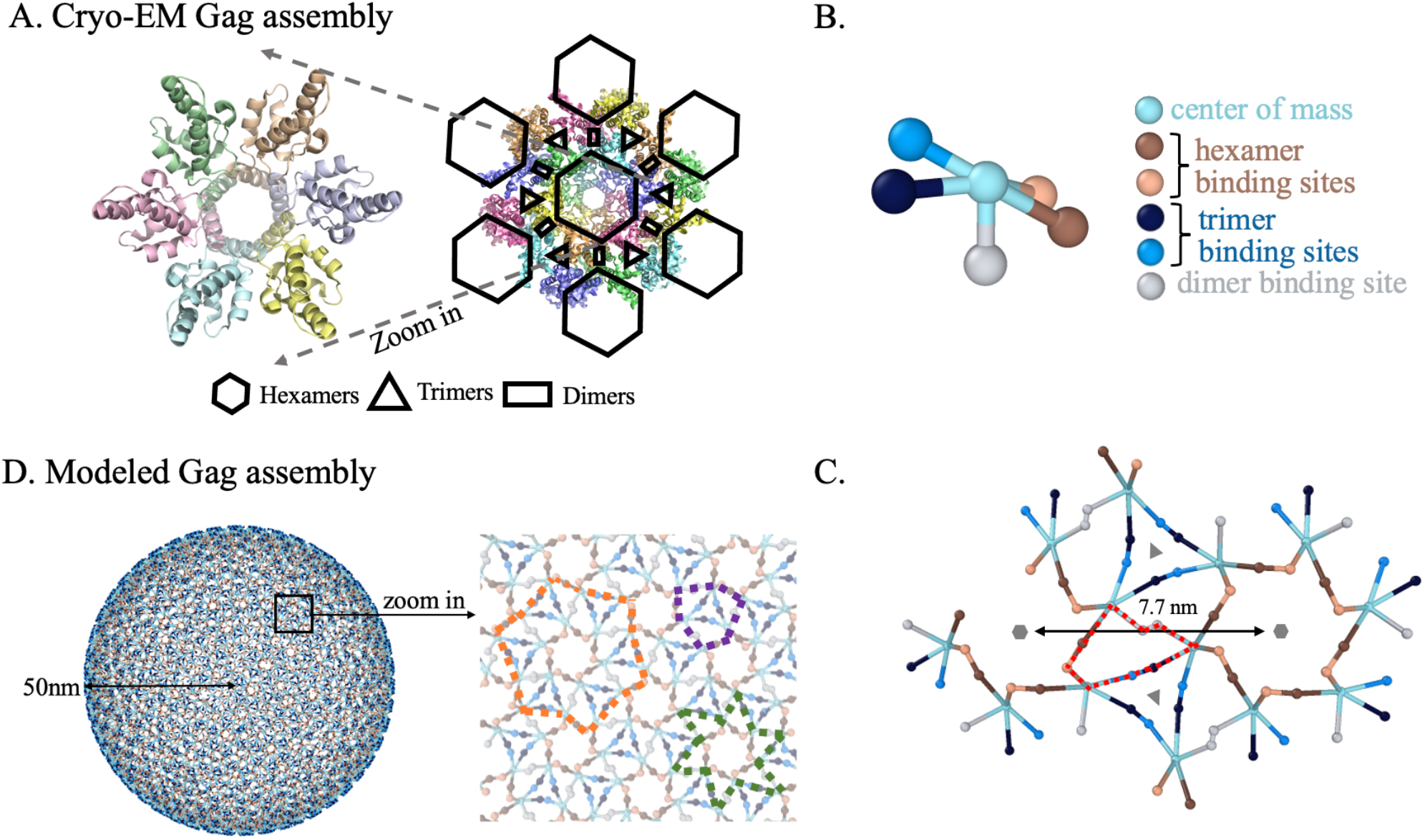
Coarse-grained model of Gag monomers and their binding interactions. (A) The cryoET immature Gag lattice from PDB ID 5I93. On the left, 6 distinct Gag monomers form a hexameric ring with each Gag contacting its two neighbors. The right image includes 18 monomers defining how the larger lattice assembles from hexagons through dimer contacts, with trimer contacts adding additional stabilizing interfaces. (B) Each coarse-grained Gag monomer has a center of mass site and 5 distinct interfaces that mediate the three types of interactions. The hexamer and trimer interactions both involve a front-to-back contact (like actin polymerization) between distinct interfaces. (C) Geometry of the coarse-grained Gag monomers assembled into the lattice, showing all three types of interactions (see Methods). In addition to the stabilizing cycles of the hexamer and trimer, cycles are also formed involving all three types of interactions (red dashed lines). (D) The spherical Gag lattice was assembled from a population of monomers, requiring ∼3700 Gags to close. Some defects in the lattice are unavoidable as a rigid hexagonal lattice cannot perfectly tile a spherical surface. From the zoomed-in structure, extra cycles formed through any two interactions can be observed. The purple line follows dimer and hexamer interactions; the orange one trimer and hexamer interactions; the green one dimer and trimer interactions.

A tri-hexagonal lattice cannot perfectly tile a spherical surface; a sphere requires pentagonal inclusions as well. However, since the hexagons are small relative to the size of the sphere, the defects that emerge due to some imperfectly aligned contacts are minimal. We allow imperfectly aligned proteins to still form bonds if they are within a cutoff distance 10% longer than the binding radius *σ*. In this way, these contacts still contribute to stabilizing the lattice, which also mimics the inherent flexibility of molecules themselves that is not captured by the rigid orientations imposed here.

### Reaction-diffusion (RD) simulations

All simulations are performed using the NERDSS (Non-Equilibrium Reaction Diffusion Self-assembly Simulator) software. NERDSS solves a particle-based and structure resolved reaction-diffusion (RD) model using the free-propagator reweighting algorithm(36). The method is derived from Smoluchowski’s model for reactive collisions between diffusing particles and has been rigorously tested to produce accurate binding kinetics and equilibria in 3D solution(37) and with the addition of rotational motion and assembly(38). The software and executable models are available open source at github.com/mjohn218/NERDSS, and all analysis described below is available as Python code at github.com/mjohn218/ioNERDSS.

We briefly outline here how the simulations work. *Step 1: Initializatio*n. Copies of molecules are placed randomly inside a rectangular box, followed by steric overlap check for all reactive binding site pairs to ensure they are not at a distance less than the binding radius *σ* for the reaction. Molecules that are titrated into the system (zeroth order reaction) are also placed randomly within the box without steric overlap. *Step 2 Reactions*. Reaction and diffusion are treated separately, and one molecule can only participate in one of them within each time step. We evaluate reactions first by calculating the reaction probability of zeroth (creation), first (dissociation), and second order (bimolecular) reactions. Dissociation reactions are modeled as Poisson processes, occurring with probability *p*_*dissoc*_ (Δ*t*) = 1 − exp (−*k*_b_Δ*t*), with intrinsic off-rate *k*_b_. All bimolecular reactions are reversible unless explicitly stated. The probability of bimolecular association events *p*_*assoc*_ (Δ*t*|*r*_0_) within timestep Δ*t* depends on the initial separation *r*_0_ of reactant sites A and B, microscopic association rate *k*_a_, the total diffusion coefficient of reactants *D*_tot_ =*D*_A_ +*D*_B_, and binding radius *σ*. The exact value is calculated from the Green’s function solution of Smoluchowski’s model, with reweighting applied due to the simple Brownian updates used for particle displacements(37). The binding probability is independent of the orientation of the two sites, it depends on only the separation. If an association reaction is accepted via comparison of the reaction probability with a uniform random number, the two molecules will be translated and rotated into the user-specified binding orientation. We reject association events between multi-protein complexes that result in steric overlap between protein monomers. We also reject association events that produce unphysically large rotational re-orientations of complexes to ‘snap’ them into the proper binding orientation. This is controlled by a scalar (scaleMaxDisplace) that multiplies the expected displacement due to translational and rotational diffusion to define a cutoff (see table 1). *Step 3 Propagation*. If the molecules within a complex do not undergo a reaction, the rigid complex diffuses both translationally and rotationally along three orthogonal axes *x, y* and *z*. Translational diffusion along any axis is simply a Brownian motion *x*(*t* + Δ*t*) = *x*(*t*) + Δ*x*, where the displacement Δ*x* follows a gaussian distribution with mean zero and standard deviation 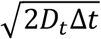. *D*_*t*_ is the translational diffusion constant. Similarly, rotational diffusion characterizes molecular rotation around a specific global axis, *θ*(*t* + Δ*t*) = *θ*(*t*) + Δ*θ*, where Δ*θ* follows Gaussian distribution with mean zero and standard deviation 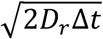. *D*_*t*_ is the rotational diffusion constant. The diffusion coefficients of complexes slow as their size increases, consistent with the assumption that the hydrodynamic radius is additive (38).

**Table 1.**
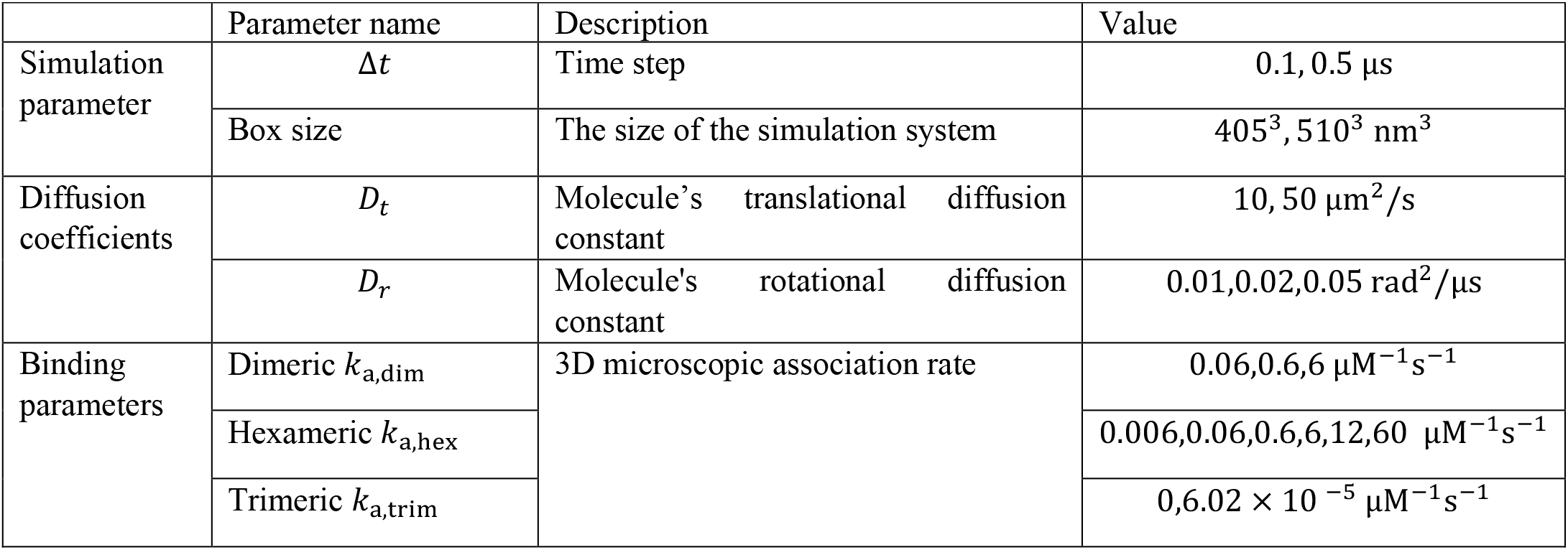
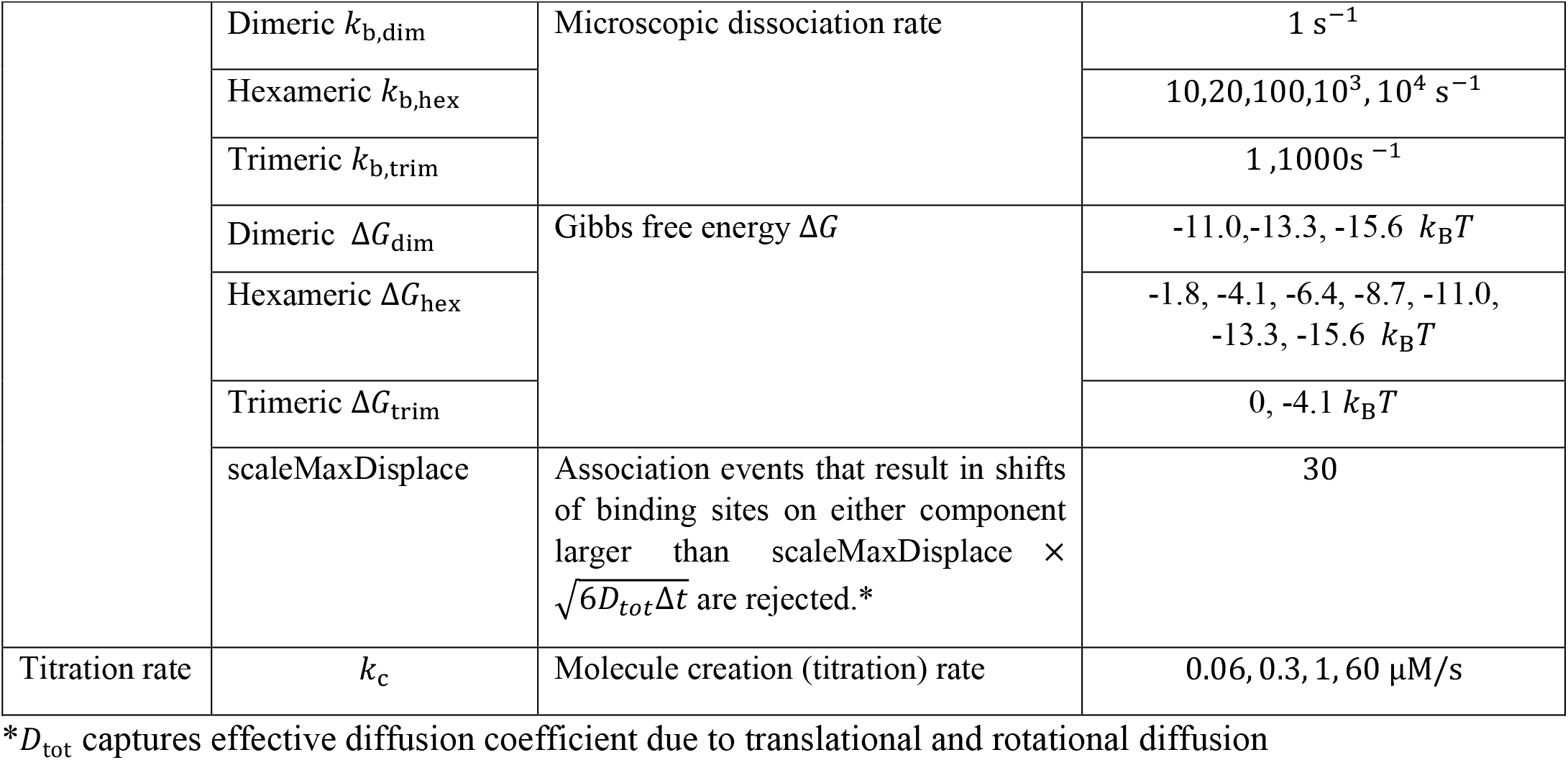
Simulation, energetic and kinetic parameters for Gag interactions.

### Simulation conditions

We perform simulations in a rectangular volume with reflective boundary conditions at the walls. Our simulations all reach a final concentration of 50 μM, consistent with experiments on immature Gag lattice assembly in solution(11). We use a larger volume of 510^3^ *nm*^3^ that contains ∼3700 Gag proteins and thus allows the formation of a complete spherical lattice. We typically use a smaller box with volume 405^3^ *nm*^3^ that contains ∼2000 Gag proteins maximally, because it is computationally more efficient and still reports on the kinetic challenges to assembling large lattices. In addition to ‘bulk’ simulations with all copies present at time zero, we also perform titration simulations where initially 10 molecules are present, and remaining proteins stochastically enter the system with rate *k*_*c*_ (see Table 1).

### Energetic and kinetic parameters

All parameters used in simulations are listed in Table 1. The stability or free energy Δ*G* of each pairwise interaction is determined by the binding (*k*_a_) and unbinding (*k*_b_) rates using standard relationships, 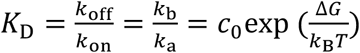, where *c*_0_ is the standard state 1M, Δ*G* = *G*_bound_ − *G*_unbound_, and *k*_on_ is the corresponding macroscopic rate defined in 3D by

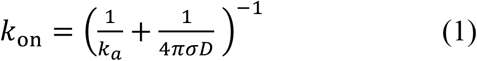

(see e.g. ref(37)).

### Fitting of association kinetics to analytical theory

To quantify how the formation of higher-order contacts impacts the assembly kinetics, we analyzed the kinetics of pairwise bond formation of both dimers and hexamers. If the bonds formed independent of any lattice, their kinetics would match up with theory from rate equations, given that our components are initially well-mixed and in 3D, which minimizes any spatial effects. These analytical solutions are known, and for completeness the kinetics of the reversible (A+A ⇌ C) reaction is given by:

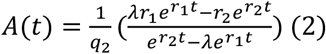

where 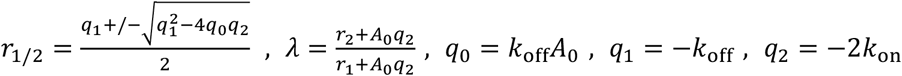. For distinct reactants, *q*_1_ and *q*_2_ change slightly (see e.g. ref(39)).

The effect of diffusion is captured in the macroscopic rate constant using Eq. 1. Given this equation, we can then fit our simulated data on bond formation, where in the case of dimer association we treat only dissociation as a fit parameter, 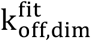. For slow hexamer interactions, we recover 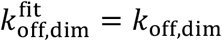, validating that the kinetics is correct as designed and dimer bonds form fully independently at short enough times. As hexamers start to form more quickly, or as trimers start to form, we see slowed dissociation of dimers, such that 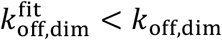. We perform the same procedure for hexamer bond-forming kinetics, except that the sites are distinct (A+B ⇌ C). For the hexamers, both the association and dissociation kinetics are influenced by dimer contacts, and thus we must treat both 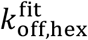 and 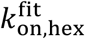 as fit parameters. We therefore then compare the ratio of these rates to their pairwise values, 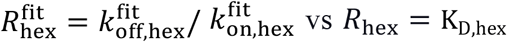. Without the dimer and trimer interactions turned on, 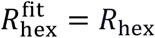 as expected, but as they are turned on, 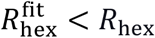, due to both faster association and slower dissociation.

### Analysis of structural regularity of assembled lattices via the Regularization Index

NERDSS records the sizes of complexes and the positions of molecules within them. To quantify how uniform and compact the growth of the Gag lattices were, we defined a Regularization Index (RI). The RI compares the surface coverage of an assembled Gag complex relative to the most compact version that contains the same number of monomers and thus has the same surface area. The most compact version is defined as growth of a spherical cap with the same radius, R=50nm. Thus, an ideal lattice that contains N_gag_ with SA_gag_=8.5nm^2^ covers a spherical cap defined by a polar angle 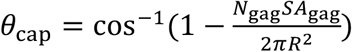. To calculate the RI, we orient our Gag lattices to be centered relative to a polar angle of zero (centered according to their center of mass) and count what fraction of the Gag monomers are enclosed within the cap defined by the deflection angle *θ*_cap_.

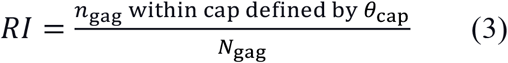

An RI of 1 thus has *n*_gag_ = *N*_gag_ and refers to ideal, compact growth. Lower values indicate more extended, fractal-like structures. See SI for further details, and note that analysis code is available open source via github.com/mjohn218/ioNERDSS

### Derivation of an optimal titration rate that avoids kinetic traps

To avoid kinetic traps, one approach is to suppress the formation of multiple competing nucleated structures while still allowing efficient growth. We design a strategy here to achieve this by keeping the concentration of Gag monomers low via a titration rate *k*_*c*_, which therefore slows nucleation. We seek to derive the expression for a rate 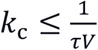, given that *τ* is the timescale for a protein to bind to a nucleated structure in a volume *V*. To quantify *τ*, consider a particle diffusing at *D*_t_ within a volume defined by the sphere of radius *R*, where the particle is initialized uniformly at any position within the volume(32). The average time for the particle to bind to a reactive sphere of radius *a* centered in that volume, given an adsorption rate of *κ* (units of length/time) is given by(32):

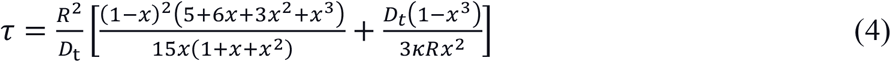

where 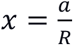. We must therefore define how our model determines the size *a* of the nucleated structure and its adsorption rate *κ*. The adsorption rate can be defined by the density of binding sites on the reactive surface, *ρ*_0_, multiplied by the association rate to bind those sites, or

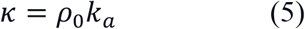

where *k*_a_ is the 3D association rate. For *k*_a_, we have three different interaction types, but because the trimer interaction is weak and does not stabilize growth, we instead can compare *k*_a,hex_ and *k*_a,dimer_. The hexamer interactions are more numerous than dimer contacts (2 per monomer), and they are more rate limiting in essentially all our models, thus we consider those binding interactions to be the time-limiting step in growth and set *k*_a_ = *k*_a,hex_. We note that if the *k*_a,dimer_ ≪ *k*_a,hex_, then we should use this rate instead and adjust the stoichiometry on the reactive sphere. The timescale *τ* scales as ∼R^3^ and thus the titration rate is most sensitive to *R* as shown in Fig S1, whereas it grows linearly with *k*_a_.

To determine the density *ρ*_0_ and size *a* of the nucleus, we must choose the number of monomers *N* in the initial nucleus and approximate this nucleus as a reactive sphere. By model construction, the length and width of each Gag monomer are approximately 4 nm and 2 nm, so the surface area taken by each Gag on the sphere is of *S*_Gag_ ≈ 8 nm^2^. If *N* is small enough, the curvature of the nucleus is minimal, and it can be represented as a disc with sticky sites on the rim, with radius 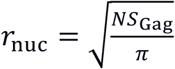. The number of binding sites on this disc is the number of free interfaces along the incomplete edge. From our model, we can estimate that the outer Gags are connected via dimer sites and that there remain free ∼0.54sites/nm. Thus, the number of sites *N*_rim_ ≈ 2*πr*_nuc_ * 0.5/nm, and we assert that the reactive sphere has radius *a* = *r*_nuc_ and binding site density 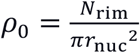. Finally, we choose a nucleus size *N* = 18, since 18 monomers are the smallest number of Gag that can assemble into a lattice containing all types of higher-order structures, i.e. the structure will contain all cycles indicated in Fig. 1D. We thus use *a* = 6.77 nm and *ρ* = 0.16 nm^−2^, and 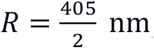, or half of the cubic box-length defining the simulation volume, and *D*_t_ = 10 μm^2^/s. When *k*_a,hex_ = 6.00 μM^−1^s^−1^, we find *k*_titr_ ≤ 3.3 × 10^−7^ M/s and when *k*_a,hex_ = 0.6 μM^−1^s^−1^, *k*_titr_ ≤ 6 × 10^−8^ M/s. Despite some approximations used to estimate these reaction parameters, they provide a well-defined estimator for the rate-limiting time-scale of binding to a single nucleation site within a volume, and we see below they correctly predict a single nucleation and growth event. Our derivation assumes that *a* and *ρ*_0_ do not change as the nucleus grows. Topologically a fragment of a Gag lattice is not a reactive sphere, as monomers attach at the edge, and it is not a volume-excluding sphere as it grows, but we consider it a reasonable approximation to use fixed estimates, particularly as *a* ≪ *R*. We also assume that the binding events are irreversible. For stable enough contacts, the unbinding rate is slow compared to growth, but we do see for the weakest contacts Δ*G*_hex_ = −6.4*k*_B_*T*, ultimately the growth speed lags behind unbinding as the lattice becomes large, and additional nuclei can form.

### Analysis of light-scattering experimental data

We analyzed time-dependent light-scattering data kindly provided from recently published work (11). We estimated a conversion from the absorbance/intensity units to concentration based on the published quantification that >95% of Gag monomers, or nearly 50*μ*M, were assembled into lattices at the end of experiments. This corresponded to 0.45 absorbance units, au. To determine the initial growth rates of Gag assembly as a function of IP6 concentration, we identified the short-time regimes where growth was approximately linear, following a lag time. We extract concentrations of IP6 given that they are initially present only in the ‘seed’ volume, whereas Gag is in both the ‘bulk’ and ‘seed’ volume, with a bulk to seed solution ratio of 65:15. For IP6=1.56

*μ*M (seed stoichiometry is Gag:RNA:IP6 = 6:2:1), we linear fit over 3-15 minutes, measuring a slope of 0.0053 au/min, using MATLAB. For IP6=9.375 *μ*M (Gag:RNA:IP6 = 6:2:6), we linear fit over 1-10 minutes, giving a slope of 0.015 au/min, and for IP6=93.75 *μ*M (Gag:RNA:IP6 = 6:2:60) we linear fit over 0.5-2.25 minutes, giving a slope of 0.0656 au/min. Using the conversion above, we thus extract titration rates of 0.0098, 0.028, and 0.12 *μ*M/s with increasing IP6. Although the stoichiometry we assume will be 6Gag:1IP6, the binding event generates 6 activated Gag so for the rate estimation the factors cancel. We predict binding rates *k*_IP-Gag_ of 126M^-1^s^-1^, 60 M^-1^s^-1^, and 26 M^-1^s^-1^ with increasing IP6, using Eq. 6 defined below. If the binding to IP6 was rate-limiting in all case, we would expect the exact same rate even as the concentration increases here by 60x. The slow-down in the rate (by ∼4x) instead indicates that the growth rate of Gag assembly is being limited by both the IP6 binding and the Gag-Gag assembly kinetics.

Lastly, given these same titration rates, we estimate the rate of binding between Gag once activated, or 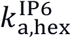. To extract this rate, we invert Eq. 8 defined below with the parameters derived from our model, solving for *k*_a_ given *k*_titr_. We show below that these model parameters work effectively in ensuring robust lattice assembly.

## RESULTS

### A. The coarse-grained Gag model can assemble a completed spherical lattice consistent with cryoET structures

To simulate assembly of the immature Gag lattice consistent with the cryo-ET structure (16), we derived a coarse-grained model of Gag monomers that form three distinct types of interactions (Fig 1). As shown in Fig 1, the hexamer and trimer interactions involve two distinct interfaces that thus allow cycles to form in a front-to-back arrangement. Homodimer interactions link together hexamers or trimers, and ultimately only two types of interactions are necessary to assemble a complete lattice. Below we test assembly both with and without trimer interactions ‘turned on’. We note that additional stabilizing cycles can form within the Gag lattice beyond just hexamers and trimers, due to the variety of intermolecular contacts between Gag, including a cycle stabilized by a trimer, hexamer and dimer bond (red dash lines in Fig 1C). Further, at larger lengthscales, additional stabilizing cycles form via a combination of any two interactions (Fig 1D). To assemble the sphere in Fig 1D from initially unbound monomers, we promoted nucleation of a single lattice by combining slow titration of monomers into a small volume with fast and irreversible binding rates. Below we characterize assembly kinetics and yield for reversible binding interactions.

### B. Kinetics of fast dimer bond formation agrees with simple theory despite higher-order assembly

By tracking the formation of dimer bonds versus time in our lattice assembly simulations (Fig 2), we find that the short-time kinetics of dimer association is in close agreement with the analytical theory of populations that can only form dimers (A+A ⇌ C), even when the hexamer association rate is as fast as the dimer rate (Fig 2A). This is because the dimer is much more stable than the short-lived hexamer bond; if we make the hexamer contacts more stable, the bonds will start to assemble more synchronously. Thus, as long as dimerization is faster and/or more stable than hexamer contacts, the formation of higher-order contacts does not significantly impact the faster kinetics of monomers forming dimers. Only at times approaching ∼1 second do we see that the lattice system forms more dimers than expected for the simple bimolecular theory, showing that the formation of hexamers helps to stabilize dimer bonds against disassembly and slow the dissociation process. The comparison between the true dimer dissociation rate (*k*_dim,off_), and an effective one 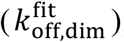, which is fit to the simulation data (see Methods), shows how this stabilization effect is stronger with faster hexamer association rate over the first second of assembly (Fig 2C). If the trimer bonds can also form, the dimers are stabilized further against dissociation over these short times, even with a slow trimerization rate of *k*_a,trim_ = 6.02 × 10^−5^ μM^−1^s^−1^ (Fig 2C). This is due to the additional cycles that are stabilized in the lattice when even weak trimer contacts can form (Fig 1). Overall, the smallest assembly unit in most of our simulations is effectively the dimer, and the kinetics is proceeding as theoretically expected for the rates we have specified. The dimer contact is known from experiments to form stably under all conditions(15) and these results are consistent with the expected higher-order lattice assembly from dimer building blocks(3).

**Figure 2.**
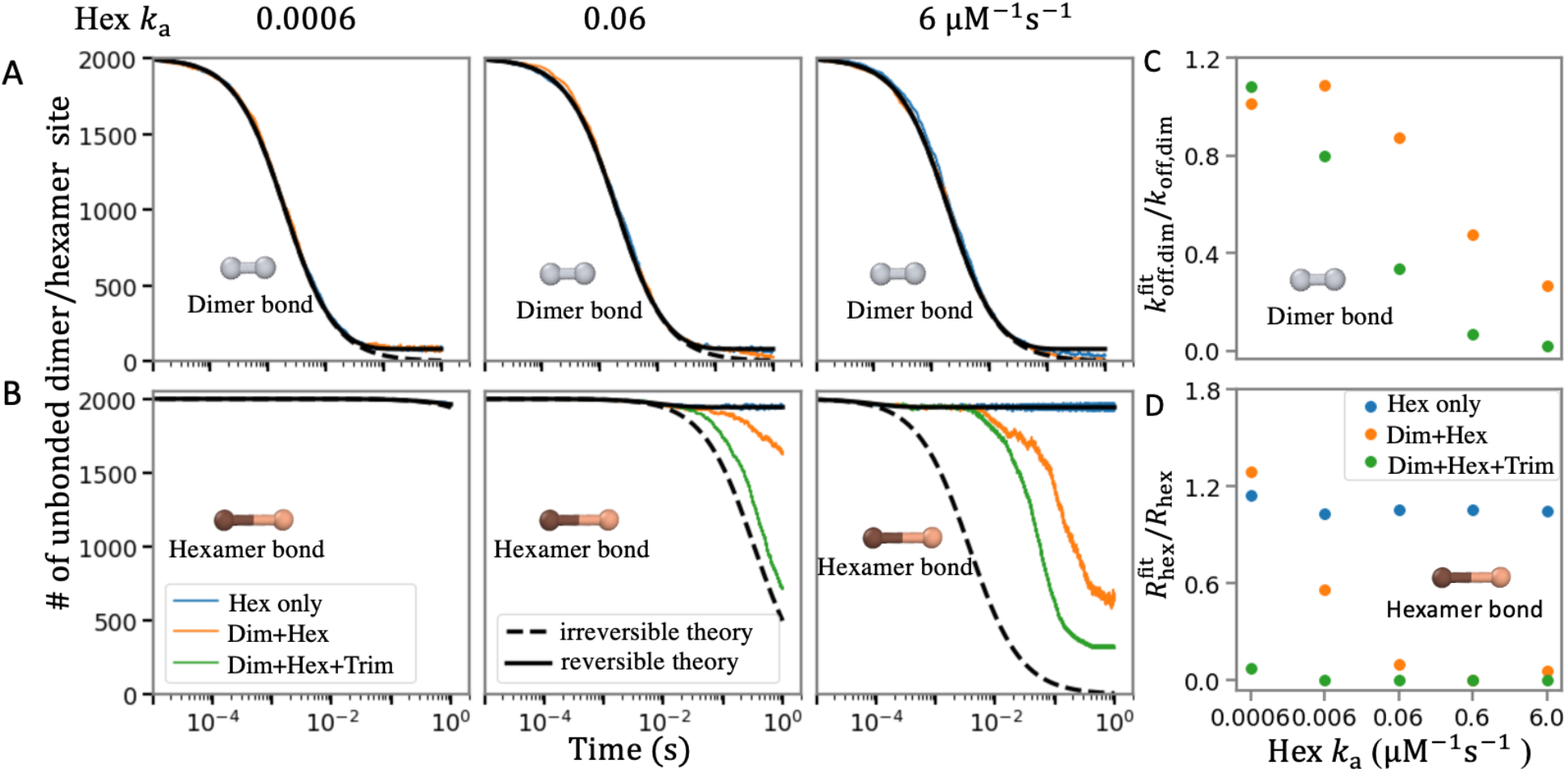
Short-term kinetics of dimer and hexamer bond formation demonstrates when higher-order assembly causes deviations from theory of bimolecular association. (A) Short-time kinetics of dimer bond formation is not sensitive to formation of hexamer and trimer interactions. The theoretical kinetics of simple irreversible (A+A→B black dashed) and reversible (A+A⇌ B solid black) association agrees with the simulations (orange and green lines), where *k*_a,dim_ = 6.02 μM^−1^s^−1^, *k*_b,dim_ = 1s^−1^. From left to right the hexamer rate is increasing, and the right-most plot has the same association rate for both hexamers and dimers, but the hexamer bonds are very short-lived compared to the dimer bonds as Δ*G*_hex_ = −6.4 *k*_B_*T* and Δ*G*_hex_ = −15.6 *k*_B_*T*. (B) The short-time kinetics of hexamer bond formation is dramatically shifted by the existence of dimer (orange) and dimer+trimer bonds (green) due to their stabilizing effect. With just hexamers forming (blue), the results match the reversible theory as expected. (C) Comparing the dimer dissociation rate *k*_dim,off_ to an effective rate that fits simulation data to theory, 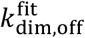 (see Methods), quantifies how this ratio 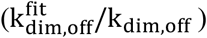 drops below 1 as the hexamer bonds start to form more quickly, slowing dimer dissociation. Δ*G*_hex_ = −6.4 *k*_B_*T* is constant for all simulations. (D) Comparing the pairwise hexamer stability to effective on and off rates fit to the data (see Methods) quantifies how 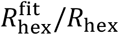 drops below 1 as dimer and trimer interactions stabilize growth. 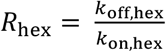. As expected, the ratio is 1 when only hexamer interactions are present (blue dots). With faster hexamer association rates, the ratio drops as dimer contacts form (orange dots) due to faster effective association and slower effective dissociation, with trimer contacts further helping (green dots) despite their relative weakness: *k*_a,trim_ = 6.02 × 10^−1^ μM^−1^s^−1^, *k*_b,trim_ = 1s^−1^, Δ*G*_trim_ = −4.1 *k*_B_*T*. All simulations have *D*_t_ = 10 μm^2^s^−1^, *D*_rot_ = 0.01 rad^2^s^−1^, Δ*t* = 0.1 μs, boxlength=405 nm.

### C. The kinetics of hexamer bond formation is accelerated by higher-order lattice assembly

Unlike the kinetics of dimer bond formation, we find that even the short-time kinetics of hexamer bond formation deviates significantly from the simple theory for independent sites (A+B ⇌ C). This is perhaps not surprising, as the hexamer bonds allow the assembly of higher-order structures that can become stabilized against disassembly. Even for the weak binding free energy of Δ*G* = −6.4 *k*_B_*T*, or *K*_*D*_ = 1,661 uM (weak relative to [C]_0_= 500 μM) we see that dissociation of the hexamer bonds slows relative to the pairwise expectations (Fig 2B). In contrast, if only hexamer bonds form (we turn off dimer and trimer interactions), the model exactly reproduces theory as expected (Fig 2). We quantified how the stabilization due to the cycles enabled by dimer and trimer interactions (see Fig 1) can effectively accelerate hexamer association and slow hexamer dissociation. We defined an effective hexamer dissociation constant, 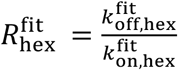, with rates defined by fitting our simulated data to the functional form of the simple theory (see Methods). As we increase the hexamer on- and off-rates while keeping the Δ*G* fixed, we see that the dissociation slows systematically, allowing faster assembly of effectively more stable hexameric contacts (Fig 2D).

These simulations also illustrate the role that the additional trimer contacts play in stabilizing the structures against disassembly. Although we add in trimer interactions at only a weak Δ*G*_trim_ = -4.1 *k*_B_*T* (*K*_*D*_ of 16.6 mM), they nonetheless quantitatively shift the kinetics of hexamer bond formation, helping to accelerate bond formation by further slowing dissociation (Fig 2D). The impact of the trimer contacts is diminished if the strength of the hexamer bond formation is increased, as we quantify further below. This is also not surprising, as stronger hexamer bonds will not need ‘help’ from other contacts to stabilize bonds. Lastly, we confirm that despite the faster formation of hexamer contacts, the equilibrium bonds formed after ∼100 seconds are the same under constant Δ*G*_hex_ (Fig S2). Adding in the trimer interaction along with the dimer ensures ∼15% more hexamer bonds form for these energies (Fig S2).

### D. Kinetic trapping emerges even for relatively weak Δ*G* due to the lattice size

By comparing systems with the same energetics but distinct hexamer binding kinetics, our systems will eventually reach the same equilibrium distribution of assemblies, but the assembly pathways they follow will be distinct. However, even for a relatively weak set of interactions (Δ*G*_hex_ = −6.4 *k*_B_*T*), we already see that our system is kinetically trapped. Specifically, for all systems, monomers and dimers have been depleted, leaving essentially no simple building blocks to complete the growth of nucleated structures (Fig.3 A,B,C and Movie S1). The size distribution of these nucleated structures varies with hexamer rates, given a fixed free energy. This shows that the pathways of assembly differ as the rates vary, forming metastable intermediates that do not converge to the same steady state over 100 seconds (Fig.3D,E,F). To grow largely complete lattices, these stalled and monomer-starved systems now must either wait for dissociation of larger complexes or wait for the rare annealing of two intermediate-sized complexes. A major factor driving trapping is the total number of monomers required to complete a lattice. Although our systems can assemble hundreds of Gag monomers into a single structure (Fig 3), the complete lattice requires 3700 monomers. For high-yield assembly, the rate of nucleating new structures should be slower than the elongation rate to complete the lattice, but adding thousands of monomers onto one structure takes significantly longer than forming a capsid that contains 10- or 100-times fewer components.

**Figure 3.**
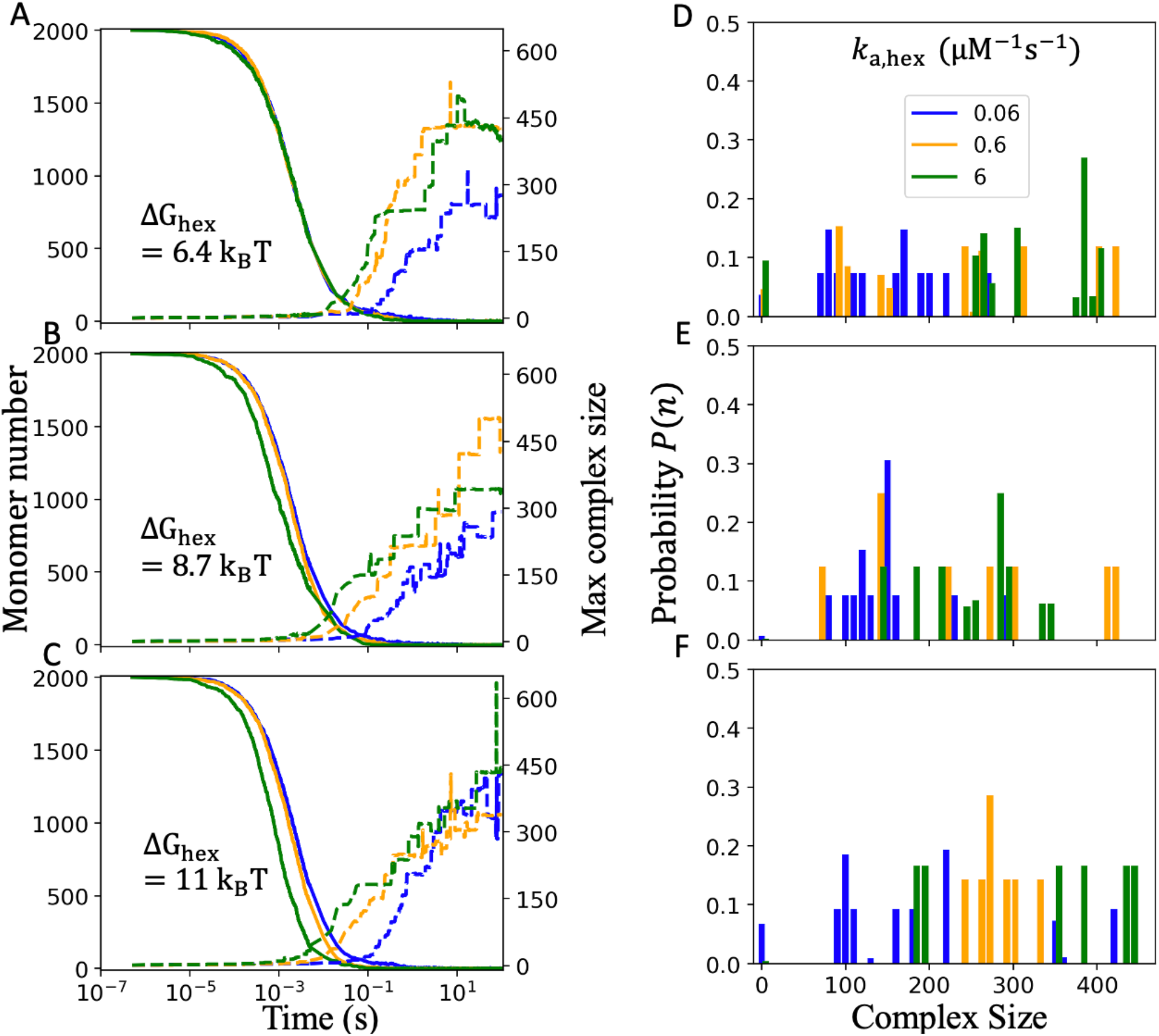
Kinetic traps emerge when all three interactions are turned on even with relatively weak ΔG. On the left column (A,B,C), we show the kinetics of monomer depletion (solid lines) and simultaneous formation of the largest assembled complex in the system (dashed lines). The hexamer rate *k*_a,dim_ Increases from 6.02 × 10^−2^ (blue), 0.602 (orange), 6.02 μM^−1^s^−1^ (green), with *k*_b,hex_ increasing to keep Δ*G*_hex_ constant. Then, from top (A) to bottom (C) the stability of the hexamer interaction ΔG_hex_ increases, by slowing *k*_b,hex_. As the small building blocks are depleted, growth stalls and the systems become trapped in incomplete intermediates (also see Movie S1). The probability distributions on the right (D,E,F) show corresponding complex sizes of intermediates present after 100 s of simulation for the same simulations to their left. All systems exhibit multiple nucleated structures, rather than a single growing structure. With faster hexamer kinetics (green), larger complexes can form on average. Even for Δ*G*_hex_ = −6.4*k*_B_*T*, only a few monomers remain free in solution, leaving no room for additional growth without dissociation. All simulations have *k*_a,trim_ = 6.02 × 10^−1^ μM^−1^s^−1^, *k*_b,trim_ = 1s^−1^, Δ*G*_trim_ = −4.1 *k*_B_*T*; *k*_a,dim_ = 6.02 μM^−1^s^−1^, *k*_b,dim_ = 1s^−1^, Δ*G*_hex_ = −15.6 *k*_B_*T*; and *D*_t_ = 10 μm^2^s^−1^, *D*_rot_ = 0.01 rad^2^s^−1^, Δ*t* = 0.5 μs, boxlength=405 nm.

As far as the assembly pathways, faster binding kinetics (both on and off rates) led to larger intermediates (Fig 3). This is consistent with the results of Fig 2, where faster hexamer kinetics combined with dimer and trimer contacts will stabilize early growth, and thus allow faster elongation of the nucleated structures before the building blocks are used up. For slower rates in contrast, more nuclei form while the concentration is relatively high, and growth is slow. This trend was preserved for the weak and stronger free energies tested (Δ*G*_hex_ = −6.4, −8.7 and -11.4 *k*_B_*T*). With all three interactions present (dimer, trimer, and hexamer), the lack of building blocks remaining after 100 seconds meant that the system was not close to equilibrium. The same trend of larger intermediates with faster association kinetics is preserved with the trimer turned on and off (Fig S5). By speeding up both the dimer and hexamer rates, we can accelerate growth, such that only 4-6 nuclei form instead of 8-10. These nucleation and growth rates are more sensitive to the hexamer association rate than to dimer and trimer rates, as we see a clear increase in the sizes of complexes formed with faster hexamerization (Fig S5).

### E. Rapidly assembled intermediates have less uniform growth

To compare the topology of our assembled structures, we defined a regularization index (RI) that measures how much a lattice deviates from a compact, ideal spherical growth (shaded with dark grey in Fig 4). Ideally, compact structures have RI=1, while a more extended fractal-like growth has increasingly lower RI. Compact structures have the shortest edge length, and thus they maximize the number of stabilizing cycles formed. Analyzing the same systems as in Fig 3, we find that smaller complexes of ∼100 Gag monomers have a higher RI on average and that the RI systematically decreases as the sizes of the complexes increases (Fig 4). This trend occurs for each Δ*G*_hex_ (Movie S1), but with less stable hexamer interactions, there is a shift toward higher RI. This indicates faster dissociation does allow structures to grow more uniformly and compactly, by correcting contacts that extend the growth and fail to satisfy as many bonds.

**Figure 4:**
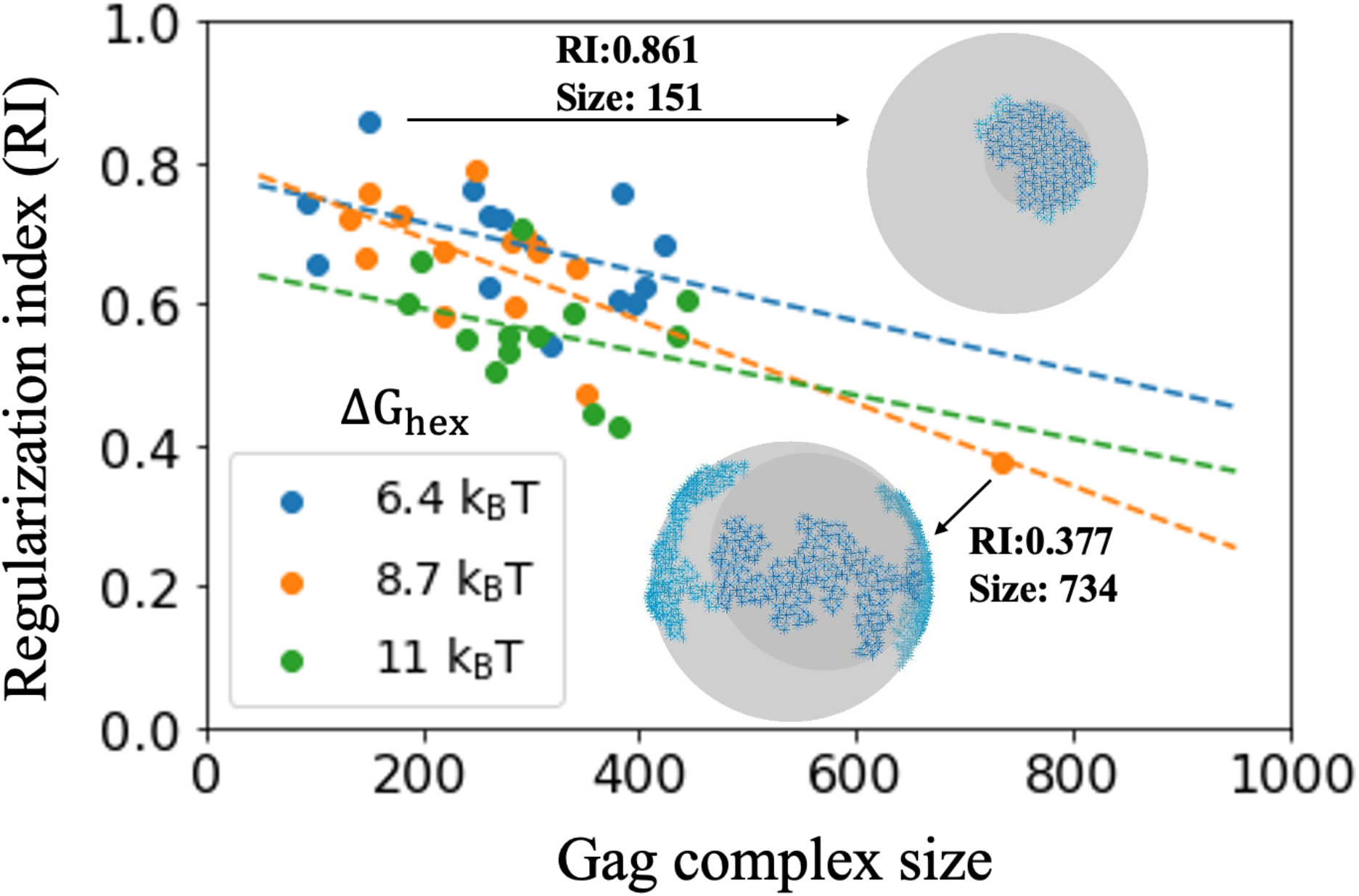
In kinetically trapped systems, growth of larger intermediates is less compact and regular. The regularization index (RI) of assembled complexes with N_gag_ >50 at steady state is on average higher for smaller assemblies. The top sphere illustrates a high RI showing significant overlap of the simulated lattice in blue with the ideal spherical cap in dark gray. The bottom sphere shows a low RI. The lattices assembled under the weaker Δ*G*_hex_ = −6.4*k*_B_*T* (blue dots) have more frequent dissociation and thus typically stabilize only with more compact structures with higher RI. Compact structures fulfill more neighbor bonds and cycles than extended structures due to their minimized edge length. The dashed lines indicate a best line fit for each Δ*G*_hex_ datapoints to highlight the trend. Same simulation parameters as Fig 3.

We further quantified growth pathways, showing they proceed primarily through addition of monomers, dimers, and small oligomers of <10 Gags, although some rare annealing events between large structures do occur (Fig S3). These annealing events are unlikely because they require that the two complexes have compatible edge geometries, and when such annealing does happen, highly irregular structures can form (Fig 4). They are also rare because the structures must be aligned properly to avoid large scale reorientation upon binding. Our algorithms will reject moves that cause any component of the assembly to move more than expected due to translational and rotational diffusion as specified by a threshold (see Methods).

### F. Retaining a dilute phase of monomers requires significantly weakened Δ*G*_hex_

Eliminating higher-order assembly requires dropping the interaction strength significantly lower than would be required for one type of interaction (Fig 5). A strong dimer interaction only reduces the total components from 50 μM of monomers to 25 μM of dimers, so we focus first on the hexamer strength, comparing Δ*G*_hex_ = −1.7, −4.1 and −6.4*k*_B_*T*. As a baseline, we consider a system that can form up to hexamers but no higher-order lattices, meaning we turn off the homodimer and trimer contacts. At Δ*G*_hex_ = −6.4*k*_B_*T*, only 0.001 % of these Gag monomers will form hexamers (Fig 5A). As we turn on interactions and increase the number of cycles possible in the lattice, the transition window from no assembly to robust assembly will shrink and shift with small changes to Δ*G*_hex_. With homodimer contacts added, a larger assembly will form at Δ*G*_hex_ = −6.4*k*_B_*T*, although >90% of components present in solution are Gag monomers, whereas at −4.1*k*_B_*T*we see no assemblies formed (Fig 5B). Finally, with all three interactions present, at Δ*G*_hex_ = −6.4*k*_B_*T*, the systems consumes nearly all monomers/dimers. Large assemblies form even with Δ*G*_hex_ = −4.1*k*_B_*T*, along with a dilute phase of monomers (Fig 5C). For this model, we return to a fully monomeric system when Δ*G*_hex_ is −1.7*k*_B_*T*.

**Figure 5.**
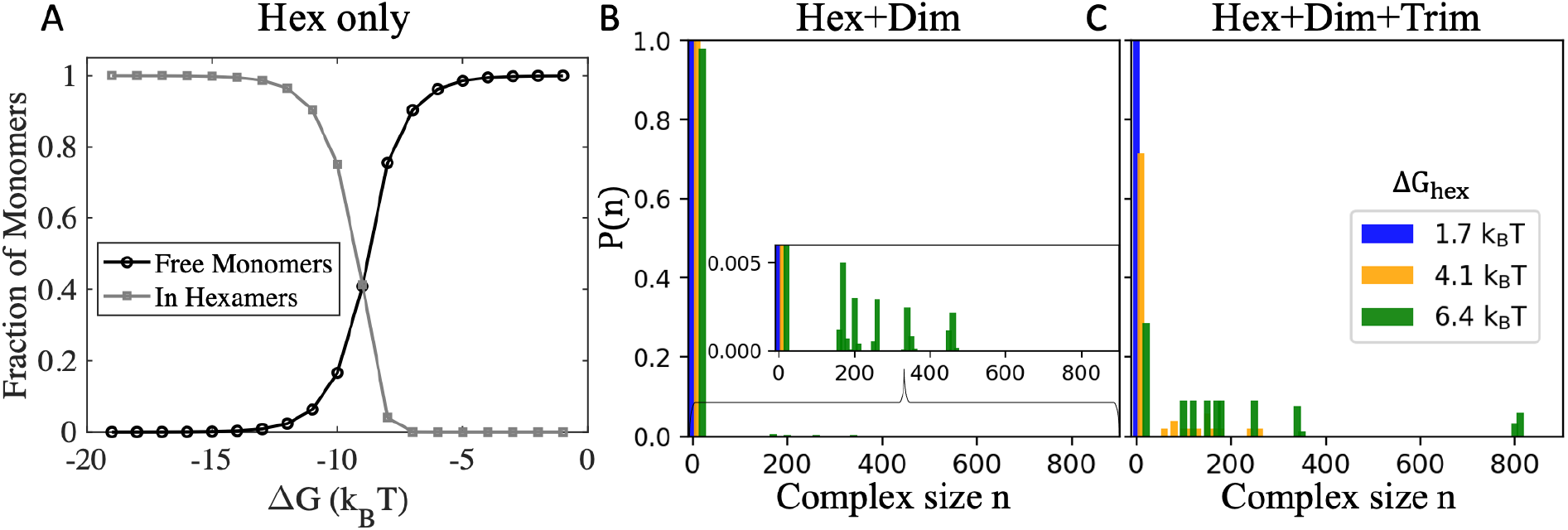
Multi-valent assembly supports lattice formation with weak hexamer contacts. A) For reference, we numerically calculated the equilibrium for Gag monomers (50 μM) that only had hexamer contacts enabled using MATLAB. The transition to <50% fraction as free monomers occurs at Δ*G*_hex_ = −8.74 *k*_B_*T*. B) For Gags that assemble the full lattice via dimer and hexamer contacts, we start to see formation of large assemblies when Δ*G*_hex_ = −6.4*k*_B_*T* (green bars), with no assembly for −1.7*k*_B_*T* (blue bars) and −4.1 *k*_B_*T* (orange bars). C) Same as (B) but now the trimer interaction is turned on at ΔG_trim_ = −4.1 k_B_T, triggering assembly at all but the lowest hexamer strength. Both (B-C) report distribution over a 5 second time window following steady-state. We note that due to the slow on-rates, some nucleation of lattices could still occur at longer times beyond those simulated here (100 seconds). The distributions are normalized over the number of distinct types of complexes, meaning that if 10 different sized complexes are present, and 1 is a monomer, than the probability of the monomer state is 0.1, even though only 1 in 2000 Gags are in the monomer form. *k*_a,dim_ = 6.02 × 10^−4^, 6.02 × 10^−3^, 6.02 × 10^−2^ μM^−1^s^−1^, *k*_b,hex_ = 1000 s^−1^, Δ*t* = 0.5 μs and *D*_t_ = 10 μm^2^s^−1^, *D*_rot_ = 0.01 rad^2^s^−1^, boxlength 405 nm, *k*_C_ = 600 μM/s. For (B-C) *k*_a,dim_ = 0.602 μM^−1^s^−1^, Δ*G*_hex_ = −13.3 *k*_B_*T*. For (C), *k*_a,trim_ = 6.02 × 10^−1^ μM^−1^s^−1^, Δ*G*_trim_ = −4.1 *k*_B_*T*.

A two-phase equilibrium with a dilute phase of monomers and dimers alongside a collection of partial assemblies is thus difficult to achieve unless only two interactions are stabilizing the lattice (Fig S4E-G). The stable dimer interaction combined with either hexameric interactions or trimer interactions at Δ*G* = −6.4 *k*_B_*T* nucleate partial assemblies, but growth ultimately stalls as the monomer concentration dilutes (Fig S4A,B). These two lattices reach the same equilibrium despite the trimer being fundamentally more stabilized against disassembly, as the higher-order cycles between dimer and trimer require more bonds than dimer and hexamer (Fig 1).

### G. Mimicking activation by co-factors can ensure slow nucleation and fast growth of a single lattice

Our simulations above demonstrate that to prevent kinetic trapping in bulk simulations for the large HIV lattice, we would need to identify the very small range of Δ*G*_hex_ values and association rates that can complete growth of the lattice before multiple nuclei form. Without the trimer, this would require slightly more stability than -6.4 *k*_B_*T* (Fig 5), but weaker than -8.7 *k*_B_*T* (Fig S4), and with faster association rates (at 50 μM). With the trimer, values would shift to between -4.1 to - 6.4*k*_B_*T*, and again with fast association. However, assembling from bulk monomers is not a problem the HIV lattice has evolved to solve. Both *in vitro* and *in vivo*, the Gag lattice assembly is only productive with co-factors, and therefore an alternate strategy for our model is to mimic the behavior of co-factors. Binding to co-factors effectively changes the concentration of ‘active’ or assembly-competent Gag monomers in a time-dependent fashion. In the bulk simulations above (Fig 3-5), the activation was therefore instantaneous, implying a high concentration of co-factors and efficient binding.

Instead of considering the dependence of activation on both the concentration and binding rate of co-factors to the Gag monomers, we simplify our approach and use a titration rate *k*_*c*_. From mass action kinetics, the rate of activated Gag production is then 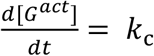. For the explicit bimolecular process, we have 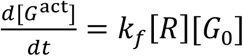, where *k* is the binding rate between Gag and co-factor, [*R*] the concentration of co-factor, and [*G*_0_] the concentration of bulk Gag prior to activation. By comparing these two we can map our titration rate to the effect of co-factor binding,

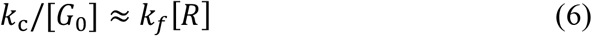

which is an accurate approximation at earlier times where the concentrations are close to their initial values. This comparison to a titration process will not directly match the time-dependence of the bimolecular association process as concentrations become limited and thus depleting with time, in which case the bimolecular process slows relative to titration. Nonetheless, it establishes useful bounds on the timescales of binding as they relate to a single titration rate: if slow titration is necessary, this would require a lower concentration of co-factor or slower binding of co-factor to Gag. In Fig 6 we test how introducing a titration process to ‘activate’ the Gag monomers can reduce the number of nuclei and thus improve the formation of more complete lattices. If the titration is too fast relative to the binding between the Gag monomers, however, improvement is minimal, indicating how this rate must be calibrated relative to the Gag assembly kinetics (Fig 6).

**Figure 6:**
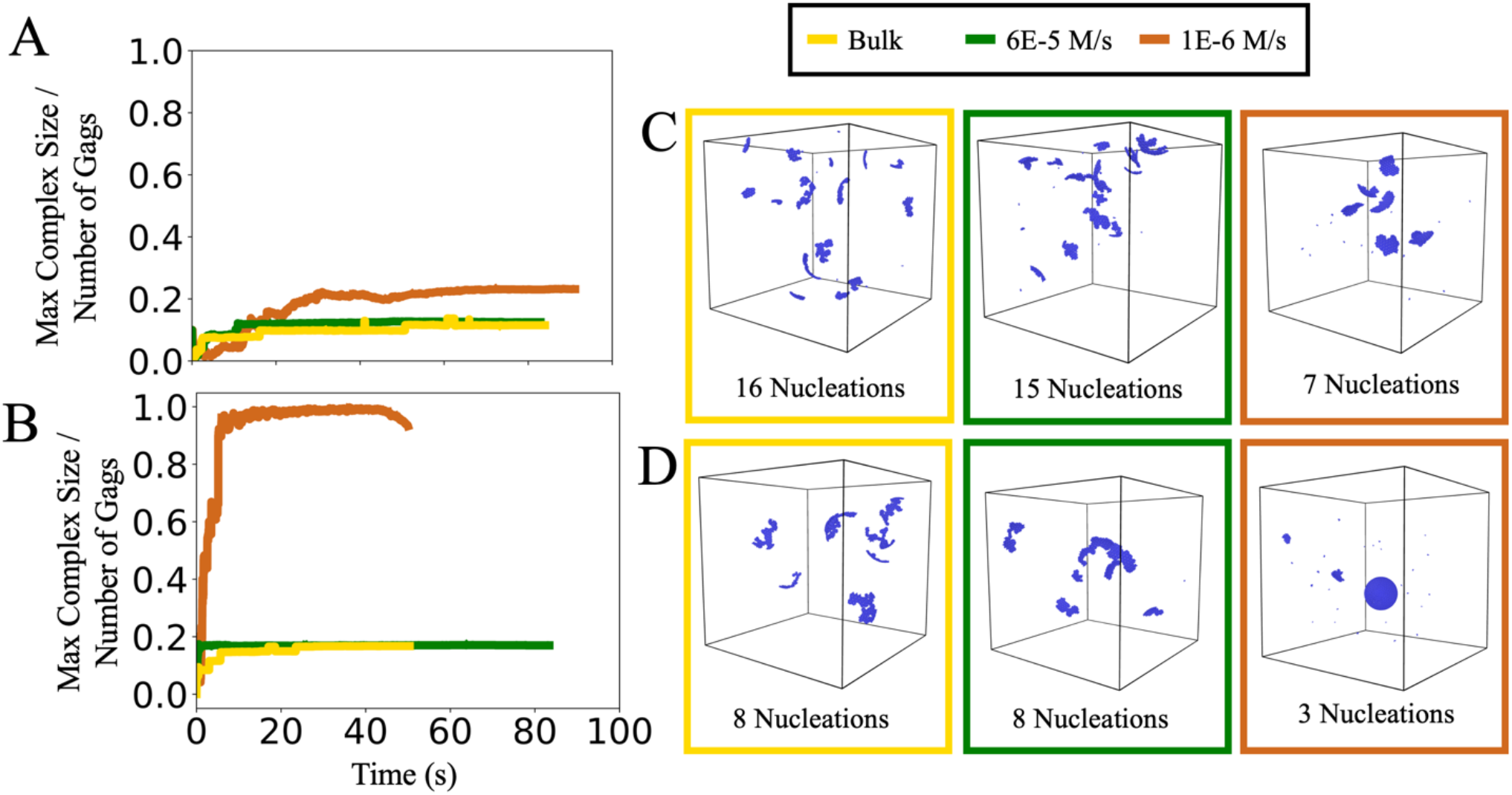
Titrating in Gag monomers can reduce nucleation events and improve lattice growth. A-B) The largest lattice formed as a function of time is normalized by the number of Gag monomers, so 1 is 100% of Gag in a single complex. The figure shows bulk simulations (yellow) and titration with 60 μM/s (green) and 1μM/s (orange). Titration was stopped when the final concentration of 50 μM was reached, which occurred after 0.83s (green) or after 50s (orange). (A) has slower binding interactions, *k*_a,dim_ = 0.06 μM^−1^s^−1^ and *k*_a,dim_ = 0.0006 μM^−1^s^−1^ *k*_b,hex_=1s^−1^. (B) has faster binding, *k*_a,dim_ = 12 μM^−1^s^−1^ and *k*_a,dim_ = 0.6μM^−1^s^−1^ *k*_b,hex_=20s^−1^. For both (A-B) *k*_b,dim_ = 1s^−1^, *k*_a,trim_ = 6.02 × 10^−1^ μM^−1^s^−1^, *k*_b,trim_ = 1s^−1^. For the slowest titration rate (orange) in (B), a single nuclei was grown during most of the simulation. In (C) and (D), we show a representative snapshot from the end of the corresponding simulations. In (B) and (D), the slowest-titration simulation uses a larger (510 nm)^3^ box to allow a lattice to complete, which it nearly does in this single trajectory (bottom right) before a few small nuclei form.

We note that the growth of our lattices does show some dependence on the diffusional search of the monomer. Simulations in larger boxes (510nm vs 405nm box length) reach the same concentrations in time, but the larger box allows the complete ∼3700 monomer lattice to form. That means that forming a single nucleus in the larger box requires the monomers to travel a larger space before ‘finding’ the nucleus. This increased travel time slows growth of the nucleated lattice and results in higher likelihood of a new lattice forming. Thus, the titration rate should reflect the volume needed to complete a lattice. We also found that slowing down the diffusion coefficients of the monomers could shift the distribution of complexes to slightly more nucleations (Fig S6). This result could not be primarily attributed to just a slower search, as a similar trend occurred when we changed the time-step. Instead, we found that our criteria for rejecting association events that involve large rotational motion by larger complexes was more permissive with faster diffusion (*D*_t_=50 μm^2^/s, *D*_r_=0.05 rad^2^/μs versus 10 μm^2^/s and 0.01 rad^2^/μs). This means that some of the early growth of the complexes is occurring due to moderately sized oligomers combining to maintain fewer nucleation sites. Hence, although the number of bonds formed is equivalent regardless of spatial extent and diffusion times, the distribution of complexes formed can skew towards more nucleation with slower diffusion.

### H. The titration rate can be derived to promote nucleation of complete lattices

From Figure 6 we clearly see that slowly titrating in or ‘activating’ Gag monomers can improve nucleation and growth by keeping the concentration low. Here we derived an optimal titration rate to ensure the growth of completed lattices. The idea is to limit the rate that a new protein appears in the simulation volume to below the rate that a protein will bind to a single nucleated structure in the volume. In this way, each new protein will contribute to growth of the single nuclei rather than formation of a distinct one. Hence, we seek to derive an expression:

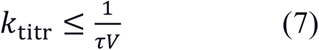

where *τ* is the timescale for a protein to bind to a nucleated structure in a volume *V*. We define the timescale *τ* by mapping to a well-defined problem in the theory of diffusion-influenced reactions for binding to a reactant in a fixed volume(32). We ignore the dissociation of proteins, or assume binding to the nuclei is irreversible, and of the two main stabilizing reactions (dimer and hexamer), we quantify binding times using the slower, or in this case, the hexamer rate. Our predicted rate is then:

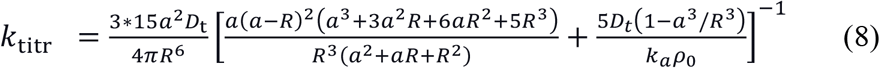

where *D*_t_ is the monomer diffusion constant, *a* ≈ 7nm is the radius of an initially nucleated structure, *k*_a_ = *k*_a,hex_ and *ρ*_0_ ≈0.16nm^-2^ is an approximate density of binding sites on a Gag lattice (see Methods for details). Lastly, what is the appropriate value for *V* in a real system, or the length scale *R* that confines our assembly process? Both *τ* and *V* scale with *R*^*3*^, hence the sensitivity of *k*_titr_ to *R* (Fig S1). The volume must be physically large enough that a complete lattice can form (with *R*_lattice_=50nm) and contain enough monomers for the complete lattice given the concentration of Gag in solution. Thus we have that

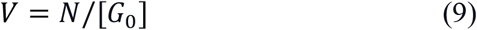

where *N* is the total monomers needed to complete a lattice (∼3700 for a 50nm sphere), and [*G*_0_] is the concentration of Gag monomers in solution. Hence, a smaller volume is needed with a higher concentration of Gag. The smaller *R* we choose, the faster the titration rate we can use, so ultimately, the rate of titration or ‘activation’ must be calibrated to the concentration of Gag in solution to form completed lattices. This lengthscale will also ultimately control the concentration of lattices in solution, [L], as assuming all monomers add to a complete lattice, [*L*] = *k*_titr_*t*_titr_/*N* where *t*_titr_ is the length of time over which the monomers are titrated to reach the target Gag concentration. We used a value of *R=*202nm for our derivation, comparable to our simulation box size. The corresponding simulations (Fig 7) then reached the same Gag concentrations as our previous ones, with 50 μM of Gag. We note that this volume does not contain the copies necessary for a complete lattice, which would instead be *R=*308nm. For the larger volume, a slower titration rate is predicted, although we see below that the assembly process can partly tolerate increases in the simulation volume.

**Figure 7:**
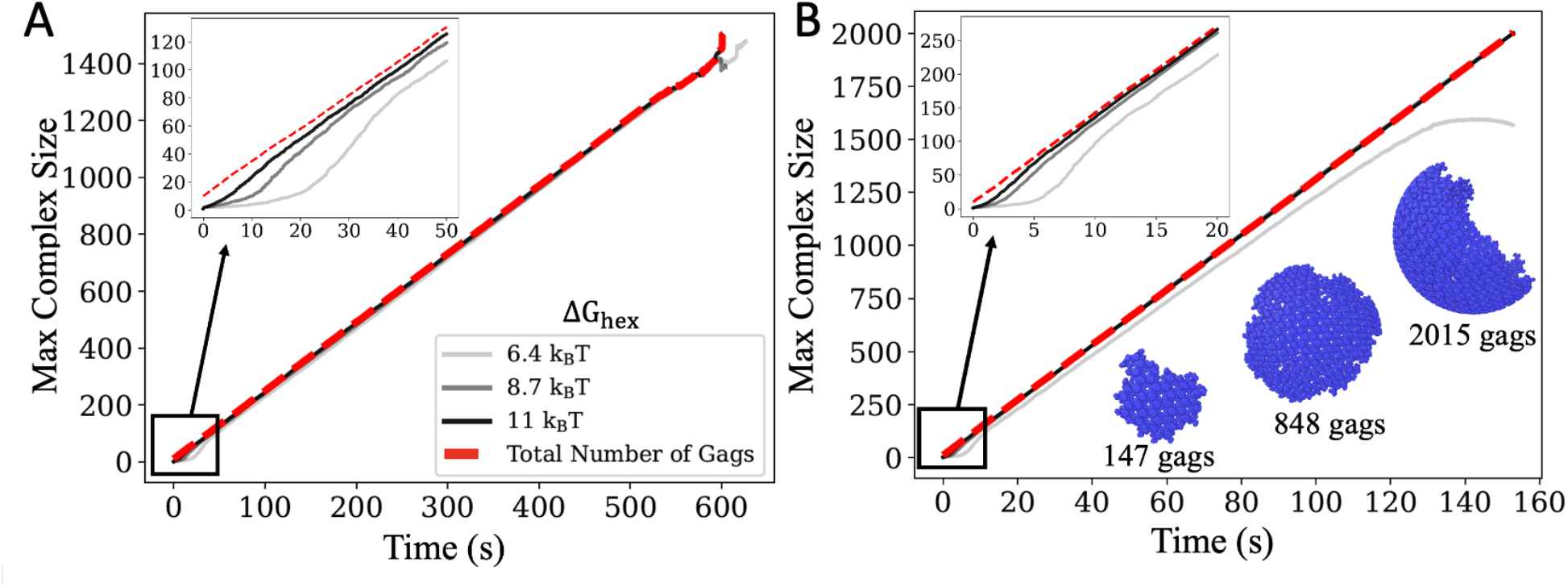
Derived titration method ensures productive nucleation and growth. (A) A single complex forms as we titrate in monomers using our derived rate *k*_titr_=0.33 *μ*M/s. All data is an average over 48 independent trajectories per model. *k*_a,dim_ =6 *μ*M^-1^s^-1^ for each curve, but Δ*G*_hex_ increases from -6.4*k*_B_*T* (light gray) -8.7 *k*_B_*T* (gray) -11 *k*_B_*T* (black) (Movie S2). The total number of Gags is shown as the dashed red line. The inset shows the initial lag time which extends with faster dissociation (light gray). (B) Here the hexamer association rate was slowed, *k*_a,dim_=0.6 *μ*M^-1^s^-1^, and *k*_titr_=0.06 *μ*M/s. Example structures from a trajectory illustrate the compact ordered growth. For all systems, *k*_a,dim_=0.6 *μ*M^-1^s^-1^, *k*_b,dim_=1s^-1^, *k*_a,trim_ = 6.02 × 10^−1^ *μ*M^-1^s^-1^, *k*_b,trim_=1s^-1^

In Figure 7 we show that this method provided excellent control to ensure the growth of a single nucleated lattice. We tested two hexamer association rates (*k*_a,dim_ = 6 and 0.6 μM^−1^s^−1^) and three hexamer stabilities (Δ*G*_hex_ = −6.4, −11.0, −13.3 *k*_*B*_*T*). The systems remain free of kinetic traps as they nucleate and grow, forming a single large assembly in our volume (Movie S2). The one exception is the weakest lattice, which after reaching a size containing ∼1200 monomers does start to nucleate additional structure in a fraction of the trajectories. The reason is that the association is counterbalanced by more frequent dissociation reactions for this weak interaction. Our simulations indicate that growth is therefore slowing somewhat as the lattice is getting larger, which would explain why ultimately new nuclei are able to form before the titrated monomers are added onto the existing structure, but they do not do so earlier in the trajectories. This further suggests that the growth of this lattice, with Δ*G*_hex_ = −6.4*k*_*B*_*T*, is not strongly stabilized against the competing disassembly, and thus assembly is less robust, requiring more tight temporal control to complete lattice formation.

Our simulations also show how the nucleation of the initial structures is dependent on both the on- and off-rates (Fig 7). The lag-time before reaching the linear growth phase is longer when off-rates are faster, consistent with more time needed to stabilize the nucleation site. For all parameter regimes these simulations produce much more compact and ideal-like growth of our structures compared to earlier simulations (Fig 3), with average values of the regularization index now ∼0.85-0.95 that are quite close to 1 at all times (ideal growth). Our derivation of *k*_titr_ above is approximate, as it assumed a fixed size and reactivity for the nucleation site, whereas in reality the nucleus grows larger and the reactive sites remain always along the edge of the lattice. Nonetheless, we find that this method keeps a low enough concentration of Gag monomers that we can now reproduce smooth and productive growth, even for highly stable lattices (Δ*G*_hex_ = −11*k*_*B*_*T*) that would otherwise rapidly become trapped into intermediates under bulk conditions (compare with Fig 5).

### I. Comparison with *in vitro* assembly kinetics reveals how slow binding of Gag to IP6 can support robust Gag assembly

The co-factor IP6 binds to Gag and activates it for assembly into the immature lattice (11, 15, 40). We therefore analyzed a recently collected set of *in vitro* experiments that tracked (via light scattering) the formation of Gag lattices vs time as a function of IP6 concentration (11) (Fig 8). Light scattering can be analyzed to quantify assembly parameters (41), but co-factors will alter the observed kinetics. Based on our theoretical arguments above, if activation of the Gag monomers was slow enough relative to the rate of Gag-Gag binding, then we would see a linear increase in Gag complex size, following a lag time. Importantly, the Gag assembly should accelerate with higher IP6 concentration if the binding of IP6 to Gag is more rate-limiting than the Gag-Gag assembly kinetics. By fitting the short-time growth in the experimental kinetics, we clearly see that with higher IP6 concentration and fixed Gag and RNA concentration, the lag-time shortens and the growth-rate increases (Fig 8A), indicating faster activation of Gag into its assembly-competent form. This short-time growth rate then corresponds to k_titr_ in our model, and if we assume the observed kinetics is limited by Gag binding to IP6, we use Eq 6 to estimate that the binding rate of IP6 to Gag is *k*_IP6-Gag_ ∼126 M^-1^s^-1^ for the lowest IP6 concentration (see Methods). Because the IP6 is initially seeded with 18% of the Gag prior to the full mixing and measurement (11), the IP6 binding could actually be slower than this rate, so we can interpret it as an upper bound.

**Figure 8.**
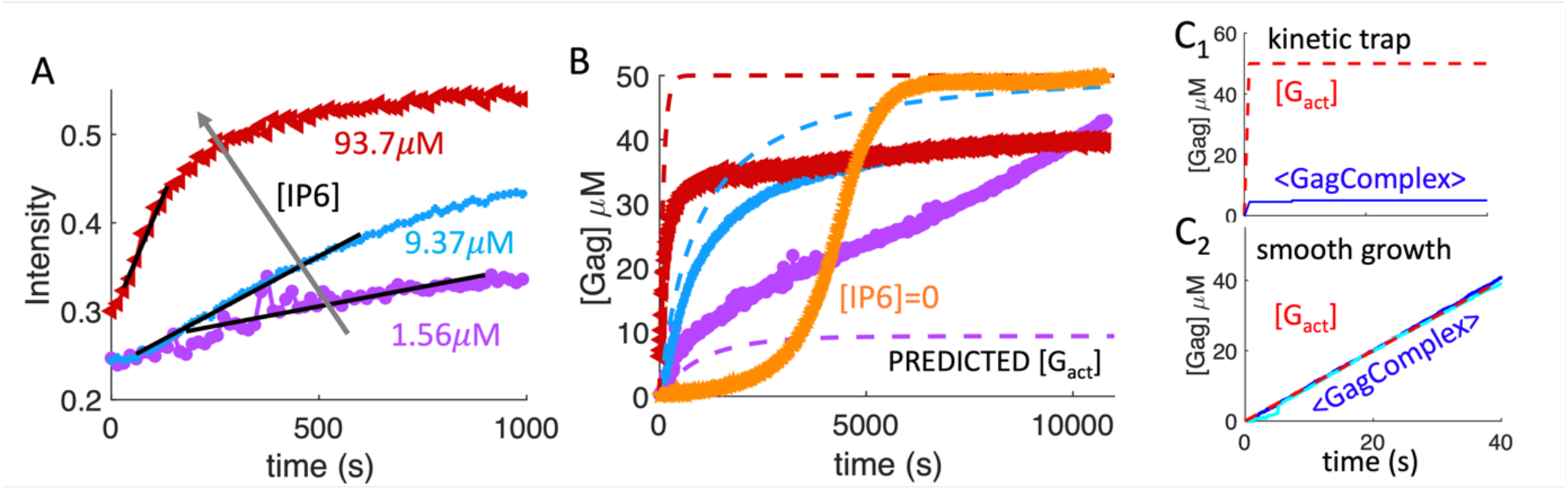
Analysis of *in vitro* Gag assembly kinetics indicates IP6 can activate Gag slowly enough to support assembly with limited trapping. A) Light scattering intensity vs time is replotted here from recent experiments measuring *in vitro* Gag Assembly (11). Gag and RNA concentration is the same for all curves, with 50 μM Gag and 3.125 μM RNA (159nt each) in solution. Concentration of IP6 increases from 1.56 μM (purple) to 9.37 (blue) and 93.7 (red), clearly driving faster Gag lattice assembly. A regime of relatively linear growth following a lag-time for each system gives a rate estimate for IP6 ‘activating’ Gag (black lines). B) Same experimental data, but we convert to Gag concentration using 0.45 absorbance units to 50 *μ*M (see Methods). We subtract 0.22 or 0.24 to re-zero the y-axis. Using our estimated rate of Gag binding to IP6, we predict the production of the IP6-activated Gag in dashed lines. This activated Gag we infer as now competent to bind other Gag much more quickly, thus allowing the Gag assembly observed experimentally. For low IP6, it is apparent that not all Gag needs to be ‘activated’ by IP6 to assemble, with a slower timescale of assembly occurring after the initial rapid assembly. This is known from the experiment: even without IP6, Gag assembly eventually occurs following a long lag time (orange). C) Our simulated data of the average size of Gag assemblies (solid blue lines, same as Fig 6B) shows the same qualitative trends given the time-dependent Gag activation (red dashed line). In C_1_) we activate the Gag too quickly compared to its assembly, and thus the assemblies become trapped. In C_2_), we activate slow enough that we see robust assembly growth over these shorter times. We see strong growth even when the box volume is larger at (510nm)^3^ (cyan data), where total copies are thus higher. We infer that the experimental results at higher IP6 (9.37 and 93.7 *μ*M) are too fast in activating Gag, leading to assembly that results in some kinetic trapping, or a plateau that does not reach >95% yield.

If the growth rate of Gag assembly is fully rate-limited by Gag-IP6 binding, that the increasing slope in Fig 8A should predict the same rate of *k*_IP6-Gag_ for increasing IP6 concentrations. Instead, we do see a modest slow-down in this predicted rate at higher IP6 concentrations (Methods). A simple interpretation is that the Gag is now being activated by IP6 a bit faster than the assembly process can keep up, so Gag activation is not fully rate-limiting. Nonetheless, the higher concentrations of G_act_ (activated Gag) driven by faster activation still drive faster assembly (Fig 8A). In Fig 8b we illustrate how the full-time evolution of the Gag assembly can be interpreted via these binding rates. Light scattering reports on the average molecular weight of solute, so it is most directly mapped to the average mass of assemblies in simulation (Fig 8C). We predict the concentration of IP6-activated Gag in Fig 8B using our value of *k*_IP6-Gag_, which then supports rapid Gag assembly. At the high IP6 concentrations (9.37 and 93.7 *μ*M), the activation is a bit faster than assembly kinetics, which leads to initially fast growth followed by some kinetic trapping, as the growth plateaus at values below the max but shows evidence of increasing at a much slower timescale (Fig 8B). This is similar to our simulation results in Fig 8C_1_. However, in the lowest IP6 concentration, for short times the growth is more like Fig8C_2_, indicating smooth growth following activation. At the lowest IP6 concentration, we can maximally activate only 9.37*μ*M Gag, or ∼20% of the total. However, assembly still proceeds beyond these times, meaning that not all Gag need to be bound to IP6 to promote assembly, but the assembly rate will be slower. This is consistent with the known assembly dynamics absent of IP6 (Fig 8B). Finally, we can then use the observed assembly kinetics compared to the activation speed to estimate the limiting rate of Gag assembly when IP6 activates, or 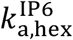.

To justify estimates on 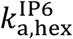, if the titration rate (IP6-driven activation rate) is too fast compared to Gag assembly, then we expect a more bulk-like condition that will promote kinetic trapping (Fig 8C_1_). If the titration rate is slow enough, the assembly kinetics will be limited by the speed of activation. Then we can estimate what the slowest assembly rate could be to ensure smooth growth, although the actual rate could also be faster; thus, we can only put a lower bound on the binding rate (Fig 8C_2_). Using the titration rate for IP6=1.56*μ*M over short times estimated as *k*_titr_∼0.0098 μM/s, we have a lower bound on 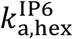 of either 7.7×10^4^ M^-1^s^-1^ using R=202nm, or 1×10^6^ M^-1^s^-1^ using R=308nm. An upper bound from the kinetics at IP6=9.73*μ*M where *k*_titr_∼0.028 *μ*M/s is then 2×10^5^ M^-1^s^-1^ using R=202nm or 3.8×10^6^ M^-1^s^-1^ using R=308nm. Hence our estimates put it between 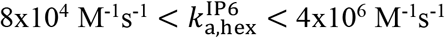. Despite the range, it indicates that the Gag-Gag assembly kinetics once activated by IP6 is significantly faster than IP6 binding to Gag (*k*_IP6-Gag_ ∼10^2^ M^-1^s^-1^), and thus IP6 can be effective in relatively slowly activating Gag for efficient assembly even when its concentrations are higher. It is also clear that Gag assembly kinetics slows when Gag must assemble lattices without all monomers benefiting from IP6 activation (Fig 8b).

## DISCUSSION

By developing a model of the immature Gag lattice from the cryoET structure, we quantified here how the strength and kinetics of the multiple Gag-Gag contacts within the lattice could support productive self-assembly from free monomers. A key component of this reaction-diffusion model is the energetic and kinetic parameters compare directly to free energies and biochemical rate constants and can thus inform pairwise interactions that are experimentally testable. A primary finding is that the Gag lattice is simply too large to assemble robustly from bulk conditions, requiring thousands of monomers to complete lattices built from monomer and dimer building blocks. Across a range of free energies and interaction rates, we showed it is remarkably difficult to complete growth of any lattice before nucleation of competing lattices, leaving the systems starved of monomers and kinetically trapped. We further show that at 50 μM of Gag monomers, suppressing assembly entirely requires very weak interactions, with K_D,Hex_ weaker than 1.6mM (−6.4*k*_*B*_*T*), or weaker than 16 mM (−4.1*k*_*B*_*T*) if trimer contacts help stabilize growth. The immature Gag lattice has not evolved to assemble from bulk components, and thus we mimicked the biological roles of co-factors RNA and IP6 to define time-dependent protocols that slowly ‘activate’ Gag monomers into assembly competent states. Our derived titration rate is quite general, applicable to a variety of self-assembling systems. We show that with this additional timescale, we can keep concentrations low and dramatically improve assembly yield for a range of Gag-Gag free energies and rates, illustrating the power of co-factors in defining both assembly kinetics and yield.

By providing theoretical justifications for our time-dependent titration protocol, we were able to analyze Gag assembly kinetics measured *in vitro* and provide here the first estimates of bounds on Gag-IP6 and Gag-Gag binding rates. Light scattering experiments showed clearly that higher concentrations of IP6 lead to faster Gag assembly kinetics(11); this observation agrees with a model of IP6 activating Gag for fast assembly. We could thus interpret the early growth rates in terms of the k_IP6-Gag_, estimating a relatively slow binding rate of ∼125M^-1^s^-1^. By also connecting to our microscopic model of titration-limited Gag assembly, we then estimated the Gag hexamer binding constant in a range of 8×10^4^-4×10^6^ M^-1^s^-1^. Thus, once Gag is activated, it can assemble relatively rapidly into the immature lattice. Though the effect of cofactors in our model is reflected in our choice of Gag-Gag binding parameters, co-factors are not explicitly modeled. Thus, we do not capture how one IP6 can stabilize formation of a 6-Gag hexamer. We simplified this stabilization as a bimolecular event with 6:1 stoichiometry in our theoretical analysis, but it most likely builds through stepwise and cooperative molecular interactions.

Including cooperativity adds more parameters that lack experimentally determined values, but our simulations are relatively efficient and can be further informed by molecular simulation. Recent coarse-grained MD simulations support that IP6 can accelerate Gag-Gag interactions, capturing coordination of one IP6 to a hexamer(33). These simulations similarly found evidence of kinetic trapping with IP6(33), consistent with our finding above that fast activation of Gag by high concentrations of IP6 results in assembly that shows signs of kinetic trapping. While it will be important in future work to explicitly capture IP6 binding and its conditional influence on Gag interactions, our results here already suggest parameters for the interactions with and without IP6 present.

For assembly of HIV immature lattices in cells, both RNA(42) and the plasma membrane(35, 43) provide additional means to temporally and conformationally control Gag activation. These are natural future extensions of our model. RNA can stimulate slower activation by introducing yet another timescale of RNA-Gag binding, which we could not quantify here because in the *in vitro* experiments, the kinetic assays start after Gag and RNA were seeded together (11). RNA-driven activation could be important for productive assembly, as cellular IP6 concentrations are high, and we showed above that if Gag is activated too quickly then it can easily become kinetically trapped. RNA can also act to tether together components, effectively localizing and concentrating components (44), with the viral genomic RNA outcompeting cellular RNA in promoting assembly (45). Membrane binding not only helps conformationally prime Gag for immature assembly(43), it also protects Gag from degradation, with >85% of the Gag produced in the cytoplasm degraded before adsorbing to the surface(26). Restriction to the plasma membrane will fundamentally concentrate components via dimensional reduction to promote assembly(46), and the localization process introduces additional timescales that can be theoretically predicted (39) to similarly ‘titrate’ the concentration. For the immature Gag lattice, the multiple sources of temporal and localized control most likely provide the robustness of assembly that is needed for such a large structure, protecting it against kinetic trapping and irregular growth.

Overall, our model simultaneously provides structural and kinetic details on the pathways of assembly as controlled by distinct Gag binding sites, further providing a means of quantifying *in vitro* kinetic measurements of Gag lattice assembly as stimulated by co-factors. Similar modeling studies of clathrin lattice nucleation and growth on membranes(47) and the immature Gag lattice dynamics following budding from the membrane using the same model here (48) demonstrate how this approach can be extended and provide quantitative connections to experimental kinetics. With this same model of the immature Gag lattice, we recently predicted (48) that activated Gag hexamer contacts would be in a range of 8-10*k*_*B*_*T*, with rates above 10^4^M^-1^s^-1^ given the more stable lattices, which is fully consistent with our model findings here. These advantages allow us to examine assembly kinetics here across the life span of immature Gag, significantly exceeding the timescales of previous computational studies of immature Gag lattices. Ultimately, simulations of self-assembly, whether using rate-based approaches (49) like here or coarse-grained energy functions (50), are critical to understand the dynamics of this nonequilibrium process that must proceed from unbound populations to functional assemblies, whether of viral capsids(18-20), immature HIV lattices(33-35) (48), or mature HIV capsids (51, 52). Coupling to cellular factors in simulations has in several cases illustrated increased robustness to assembly(29, 35, 47), with modeling approaches thus offering both general and system specific guides to interpret and integrate with further quantitative experiments.

## SUPPLEMENTAL MOVIES

**Movie S1:** Assembly kinetics with 50 μM bulk Gag shows kinetic trapping. Δ*G*_hex_ = −11.00 *k*_*B*_*T k*_a,dim_ = 6 μM^−1^s^−1^.

**Movie S2**. Assembly kinetics with *k*_titr_ = 0.33 μM/s allows for slower Gag activation and robust and compact assembly. Δ*G*_hex_ = −11.0 *k*_*B*_*T k*_a,dim_ = 6 μM^−1^s^−1^.

## ACKNOWLEDGEMENTS

MEJ gratefully acknowledges funding from an NSF CAREER Award 1753174. Our work used the rockfish cluster of the ARCH supercomputing system at Johns Hopkins University, supported by NSF MRI 1920103. We are grateful to Prof. Owen Pornillos for kindly providing his data and discussing the experimental conditions.

## Notes

### Competing Interest Statement

The authors have declared no competing interest.

## REFERENCES

1. Freed, E.O. and A.J. Mouland, The cell biology of HIV-1 and other retroviruses. Retrovirology, 2006. 3: p. 77.

2. Schur, F.K., et al., Structure of the immature HIV-1 capsid in intact virus particles at 8.8 A resolution. Nature, 2015. 517(7535): p. 505–8.

3. Tan, A., et al., Immature HIV-1 assembles from Gag dimers leaving partial hexamers at lattice edges as potential substrates for proteolytic maturation. Proc Natl Acad Sci U S A, 2021. 118(3).

4. Pettit, S.C., et al., Initial cleavage of the human immunodeficiency virus type 1 GagPol precursor by its activated protease occurs by an intramolecular mechanism. J Virol, 2004. 78(16): p. 8477–85.

5. Sundquist, W.I. and H.G. Krausslich, HIV-1 assembly, budding, and maturation. Cold Spring Harb Perspect Med, 2012. 2(7): p. a006924.

6. Mallery, D.L., et al., Cellular IP6 Levels Limit HIV Production while Viruses that Cannot Efficiently Package IP6 Are Attenuated for Infection and Replication. Cell Rep, 2019. 29(12): p. 3983–3996 e4.

7. Mallery, D.L., et al., A stable immature lattice packages IP6 for HIV capsid maturation. Sci Adv, 2021. 7(11).

8. Keller, P.W., et al., HIV-1 maturation inhibitor bevirimat stabilizes the immature Gag lattice. J Virol, 2011. 85(4): p. 1420–8.

9. Waheed, A.A. and E.O. Freed, HIV type 1 Gag as a target for antiviral therapy. AIDS Res Hum Retroviruses, 2012. 28(1): p. 54–75.

10. Bush, D.L. and V.M. Vogt, In Vitro Assembly of Retroviruses. Annu Rev Virol, 2014. 1(1): p. 561–80.

11. Kucharska, I., et al., Biochemical Reconstitution of HIV-1 Assembly and Maturation. J Virol, 2020. 94(5).

12. Campbell, S. and V.M. Vogt, In vitro assembly of virus-like particles with Rous sarcoma virus Gag deletion mutants: identification of the p10 domain as a morphological determinant in the formation of spherical particles. J Virol, 1997. 71(6): p. 4425–35.

13. Campbell, S., et al., Modulation of HIV-like particle assembly in vitro by inositol phosphates. Proc Natl Acad Sci U S A, 2001. 98(19): p. 10875–9.

14. Wagner, J.M., et al., Crystal structure of an HIV assembly and maturation switch. Elife, 2016. 5.

15. Datta, S.A., et al., Interactions between HIV-1 Gag molecules in solution: an inositol phosphate-mediated switch. J Mol Biol, 2007. 365(3): p. 799–811.

16. Schur, F.K., et al., An atomic model of HIV-1 capsid-SP1 reveals structures regulating assembly and maturation. Science, 2016. 353(6298): p. 506–8.

17. Zlotnick, A., To build a virus capsid. An equilibrium model of the self assembly of polyhedral protein complexes. J Mol Biol, 1994. 241(1): p. 59–67.

18. Zlotnick, A., et al., A theoretical model successfully identifies features of hepatitis B virus capsid assembly. Biochemistry, 1999. 38(44): p. 14644–52.

19. Hagan, M.F., Modeling Viral Capsid Assembly. Adv Chem Phys, 2014. 155: p. 1–68.

20. Mohajerani, F., et al., Multiscale Modeling of Hepatitis B Virus Capsid Assembly and Its Dimorphism. ACS Nano, 2022. 16(9): p. 13845–13859.

21. Hagan, M.F. and O.M. Elrad, Understanding the concentration dependence of viral capsid assembly kinetics--the origin of the lag time and identifying the critical nucleus size. Biophys J, 2010. 98(6): p. 1065–74.

22. Mohajerani, F., et al., Mechanisms of Scaffold-Mediated Microcompartment Assembly and Size Control. ACS Nano, 2021. 15(3): p. 4197–4212.

23. Liu, Y. and X. Zou, A New Model System for Exploring Assembly Mechanisms of the HIV-1 Immature Capsid In Vivo. Bull Math Biol, 2019. 81(5): p. 1506–1526.

24. Gartner, F.M., I.R. Graf, and E. Frey, The time complexity of self-assembly. Proc Natl Acad Sci U S A, 2022. 119(4).

25. Hagan, M.F., O.M. Elrad, and R.L. Jack, Mechanisms of kinetic trapping in self-assembly and phase transformation. J Chem Phys, 2011. 135(10): p. 104115.

26. Tritel, M. and M.D. Resh, Kinetic analysis of human immunodeficiency virus type 1 assembly reveals the presence of sequential intermediates. J Virol, 2000. 74(13): p. 5845–55.

27. Baschek, J.E., R.K. HC, and U.S. Schwarz, Stochastic dynamics of virus capsid formation: direct versus hierarchical self-assembly. BMC Biophys, 2012. 5: p. 22.

28. Boettcher, M.A., H.C. Klein, and U.S. Schwarz, Role of dynamic capsomere supply for viral capsid self-assembly. Phys Biol, 2015. 12(1): p. 016014.

29. Lazaro, G.R. and M.F. Hagan, Allosteric Control of Icosahedral Capsid Assembly. J Phys Chem B, 2016. 120(26): p. 6306–18.

30. von Smoluchowski, M., Attempt to derive a mathematical theory of coagulation kinetics in colloidal solutions. Z. Phys. Chem, 1917. 92: p. 129.

31. Rice, S.A., Diffusion Limited Reactions. Comprehensive Chemical Kinetics. Vol. 25. 1985, Netherlands: Elsevier Science and Technology.

32. Szabo, A., K. Schulten, and Z. Schulten, 1st Passage Time Approach to Diffusion Controlled Reactions. Journal of Chemical Physics, 1980. 72(8): p. 4350–4357.

33. Pak, A.J., et al., Inositol Hexakisphosphate (IP6) Accelerates Immature HIV-1 Gag Protein Assembly toward Kinetically Trapped Morphologies. J Am Chem Soc, 2022. 144(23): p. 10417–10428.

34. Ayton, G.S. and G.A. Voth, Multiscale computer simulation of the immature HIV-1 virion. Biophys J, 2010. 99(9): p. 2757–65.

35. Pak, A.J., et al., Immature HIV-1 lattice assembly dynamics are regulated by scaffolding from nucleic acid and the plasma membrane. Proc Natl Acad Sci U S A, 2017. 114(47): p. E10056–E10065.

36. Varga, M.J., et al., NERDSS: A Nonequilibrium Simulator for Multibody Self-Assembly at the Cellular Scale. Biophysical Journal, 2020. 118(12): p. 3026–3040.

37. Johnson, M.E. and G. Hummer, Free-Propagator Reweighting Integrator for Single-Particle Dynamics in Reaction-Diffusion Models of Heterogeneous Protein-Protein Interaction Systems. Physical Review X, 2014. 4(3).

38. Johnson, M.E., Modeling the Self-Assembly of Protein Complexes through a Rigid-Body Rotational Reaction-Diffusion Algorithm. J Phys Chem B, 2018. 122(49): p. 11771–11783.

39. Mishra, B. and M.E. Johnson, Speed limits of protein assembly with reversible membrane localization. J Chem Phys, 2021. 154: p. 194101.

40. Dick, R.A., et al., Inositol phosphates are assembly co-factors for HIV-1. Nature, 2018. 560(7719): p. 509–512.

41. Endres, D. and A. Zlotnick, Model-based analysis of assembly kinetics for virus capsids or other spherical polymers. Biophys J, 2002. 83(2): p. 1217–30.

42. Rein, A., et al., Diverse interactions of retroviral Gag proteins with RNAs. Trends Biochem Sci, 2011. 36(7): p. 373–80.

43. Datta, S.A., et al., HIV-1 Gag extension: conformational changes require simultaneous interaction with membrane and nucleic acid. J Mol Biol, 2011. 406(2): p. 205–14.

44. Jouvenet, N., S.M. Simon, and P.D. Bieniasz, Imaging the interaction of HIV-1 genomes and Gag during assembly of individual viral particles. Proc Natl Acad Sci U S A, 2009. 106(45): p. 19114–9.

45. Comas-Garcia, M., et al., Dissection of specific binding of HIV-1 Gag to the ‘packaging signal’ in viral RNA. Elife, 2017. 6.

46. Yogurtcu, O.N. and M.E. Johnson, Cytosolic proteins can exploit membrane localization to trigger functional assembly. PLoS Comput Biol, 2018. 14(3): p. e1006031.

47. Guo, S.-K., A.J. Sodt, and M.E. Johnson, Large self-assembled clathrin lattices spontaneously disassemble without sufficient adaptor proteins. PLOS Computational Biology, 2022. 18(3): p. e1009969.

48. Guo, S., et al., Defects in the HIV immature lattice support essential lattice remodeling within budded virions. bioRxiv, 2022. https://www.biorxiv.org/content/10.1101/2022.11.21.517392v1.

49. Sweeney, B., T. Zhang, and R. Schwartz, Exploring the parameter space of complex self-assembly through virus capsid models. Biophys J, 2008. 94(3): p. 772–83.

50. Hagan, M.F. and D. Chandler, Dynamic pathways for viral capsid assembly. Biophys J, 2006. 91(1): p. 42–54.

51. Grime, J.M. and G.A. Voth, Early stages of the HIV-1 capsid protein lattice formation. Biophys J, 2012. 103(8): p. 1774–83.

52. Grime, J.M., et al., Coarse-grained simulation reveals key features of HIV-1 capsid self-assembly. Nat Commun, 2016. 7: p. 11568.

