## Supplemental Information for "Temporal control by co-factors prevents kinetic trapping in retroviral Gag lattice assembly"

for

†co-first authors

**Supplemental Methods:**

*S1. Constructing a Gag model that generates a single target spherical curvature*

A rigid coarse-grained (CG) model is derived from the structural coordinates of the immature HIV Gag lattice in 5I93.pdb, which provides all atom positions of 18 Gags. By defining interfaces given a distance cutoff of 3.5A between residues in adjacent Gag monomers, each Gag monomer has five binding sites, including one dimer, two hexamer, and two trimer sites, which is consistent with the known stabilizing contacts. The coordinates of each interface in our model are defined by identifying all residues that participate in each interface (using the cutoff), and then averaging the position of the residue $\alpha$ Carbons to create a single site per interface. Each monomer finally contains one center of mass (COM) and five binding sites (see Illustration S1).

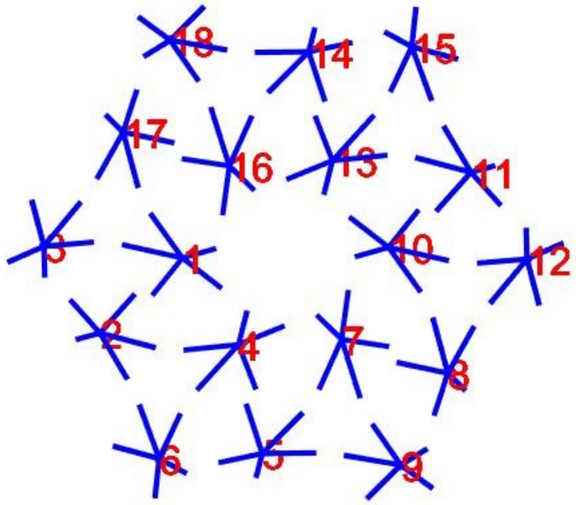


*Illustration S1. Coarse-grained rigid model of 18 Gag monomers arranged in the immature lattice from 5L93.pdb*

Each of these 3 binding interactions (dimer, hexamer, trimer) must have (up to) 5 angles associated with them to control the orientation of the monomers in the bound state.

However, these 18 Gags are not perfectly identical, and their COMs are not located precisely on one spherical surface. Thus, the angles and orientations between each pair of Gag are not perfectly symmetric. Simple averaging across all pairs does not work; the order of assembly proceeding from hexameric, dimeric, or trimeric contacts can produce inconsistent curvatures of the growing lattice. To avoid such significant defects and reconstruct a single-curvature sphere, we adjust the structure and positions of the 18 Gags to regularize the global curvature given one single set of interface coordinates, three binding lengths σ, and three sets of binding angles as required by NERDSS.

As stated above, not all 18 Gags are on the surface of the same sphere. We first need to find the "sphere of best fit" by minimizing the Root-Mean-Square Deviation (RMSD) value:

$$\mathrm{RMSD} = \sqrt{\frac{1}{18}\sum_{i=1}^{18}\left(\sqrt{(x_i - x_0)^2 + (y_i - y_0)^2 + (z_i - z_0)^2} - R\right)^2} \qquad \text{(S1)}$$

where $(x_i, y_i, z_i)$ is the COM of Gag i, $(x_0, y_0, z_0)$ is the center of the desired sphere, and R is its radius. The values of $(x_0, y_0, z_0)$ and $r_0$ should produce the smallest RMSD, which can be obtained by solving equations (2a)-(2c) :

$$\frac{\partial(\mathrm{RMSD})}{\partial x_0} = 0 \qquad \text{(S2a)}$$

$$\frac{\partial(\mathrm{RMSD})}{\partial y_0} = 0 \qquad \text{(S2b)}$$

$$\frac{\partial(\mathrm{RMSD})}{\partial z_0} = 0 \qquad \text{(S2c)}$$

To position 18 Gags symmetrically on the desired sphere calculated above, we first adjust the position of the 6 Gags in the middle to be a perfect hexamer through rotation and then determine the peripheral Gag positions by translation. First, a template “Gag” is produced by manually putting the COM of one Gag on the desired sphere and reestablishing the rigid structure (5 binding sites) around the new COM. Then we rotate it for 60˚, 120˚, 180˚, 240˚ and 300˚ around the axis passing through the sphere-center and hexagon-center, to get 6 Gags with the exact same structure forming a hexagon and all of them are on the desired sphere. For example, Gag 16 can be reproduced by rotating Gag1 for 60˚ (Illustration S2).

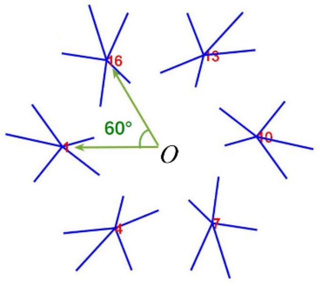


*Illustration S2. CG Model for the 6 central Gags forming a perfect hexamer on the sphere surface, with O the center of the hexamer.*

Peripheral Gags do not contribute to the central hexagon, but they are part of neighboring hexagon structures and contain binding information of dimer and trimer interactions. By translating the central hexagon on the sphere surface, we can determine the position of surrounding hexagons and obtain the position of peripheral Gags. For example, Gag5 and Gag6 are in the same hexagon and their positions are determined by Gag16 and Gag13 respectively in the central hexagon.

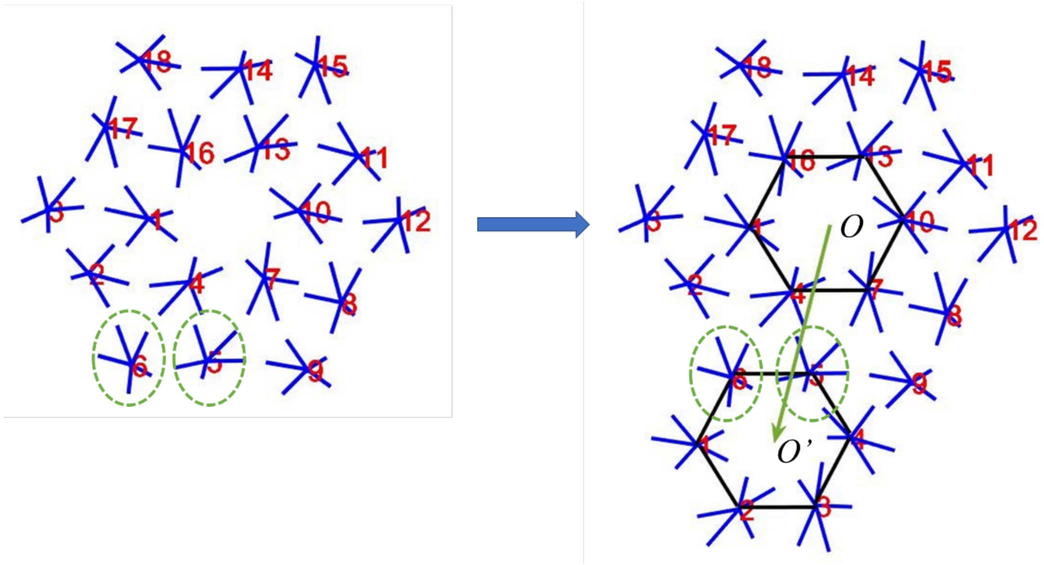


*Illustration S3. Translation of the central hexagon patterns the complete lattice.*

We can move the central hexagon as shown above. To make it simple, we suppose that the translation vector is $\overrightarrow{OO'} = \overrightarrow{r_6} - \overrightarrow{r_{16}}$, where $\overrightarrow{r_6}$ is the COM of Gag6 and $\overrightarrow{r_{16}}$ is the COM of Gag16. If we want to change the distance between hexagons, we can simply change the length of $\overrightarrow{OO'}$, but keep the direction the same. Then we move the central hexagon along the vector direction of $\overrightarrow{OO'}$ until the center of the hexagon $O$ arrives at the point $O'$. One thing to note is that the translation is carried out on the sphere surface, which is different from regular 3D translation. The detailed steps are described in the SI of previous work [1], where the diffusion of multi-site complexes on a sphere surface is discussed. Repeat the above process for every hexagon around the central one to get the position for all 12 peripheral Gags.

This symmetric structure maintains the global structure of the original one and can assemble into a complete sphere with a single curvature. Some defects eventually occur in the packing of the monomers during dynamic assembly because a hexameric lattice does not perfectly tile a sphere surface. We calculate the binding lengths σ and five binding angles from this symmetrized Gag structure based on equations presented in previous work [1]. The python code for performing this symmetrization given the PDB coordinates is available open source in the ioNERDSS python package.

*S2. Data analysis methods*

Suppose in a certain trajectory, the time step is $\Delta t$, there are total M steps, the initial time is always 0 s, and the final time is $t_f = M\Delta t$. Let $N_0$ be the number of Gag proteins in the system, so there are $N_0$ possible Gag complex sizes. Define sequence $N_i = \{n_{ij}\}$ , $i \in \{1,2,3,\dots,M\}$, $j \in \{1,2,3,\dots,N_0\}$ and $n_{ij} \in \mathbb{N}$ represents the number lattices consisting of j Gag monomers in the system at time step i. For example, $n_{10,22} = 7$ means at the 10th step, there are seven 22-mer Gag complexes in the system.

The maximum complex size: The maximum complex size of Gag at step i, $N_{i,Max}$, is defined by Eq. S3:

$$N_{i,Max} = \max\left(\arg_{X \in N_i, X>0} X\right) \qquad \text{(S3)}$$

The mean complex size: The mean complex size of Gag at step i, $N_{i,Mean}$, is defined by Eq. S4:

$$N_{i,Mean} = \frac{\sum_{j=2}^{N_0} n_{ij} \times j}{\sum_{j=2}^{N_0} n_{ij}} \qquad \text{(S4)}$$

To reduce the fluctuation of the statistics, monomers are excluded from the calculation.

The number of nucleation sites: The nucleation site is defined to be complexes that include more than 50 Gag proteins. The number of nucleation sites at step i, $Nuc_i$, is defined by Eq. S5:

$$Nuc_i = \sum_{j=51}^{N_0} n_{ij} \qquad \text{(S5)}$$

Probability of occurrence in the steady state: Suppose we know during the period $[T_{eq1}\Delta t, T_{eq2}\Delta t]$, $T_{eq1}, T_{eq2} \in \mathbb{N}, 0 \le T_{eq1} < T_{eq2} \le M$, the system is in the steady state. The probability of occurrence for j-mer in this period, $P_j$, can be calculated by Eq. S6:

$$P_j = \frac{\sum_{i=T_{eq1}}^{T_{eq2}} n_{ij}}{\sum_{k=1}^{N_0} \sum_{i=T_{eq1}}^{T_{eq2}} n_{ik}} \qquad \text{(S6)}$$

Regularization index (RI): RI measures the regularity or compactness of a Gag assembly by measuring how close the lattice is to a perfect spherical cap. Assume we wish to calculate the RI of a complex that includes J monomers. Set $P = \{p_1, p_2, \ldots, p_J\}$, where $p_i = \{x_i, y_i, z_i\}, i \in \{1,2, \ldots, J\}$ represents the coordinate of the center of mass (COM) of each Gag molecule. Each complex is compared with an equal surface area, standard spherical cap $C_{ideal}$ placed on the complex center. To calculate the RI, we follow the steps below,

1. Determine the center of the sphere that contains all points in P.

Let $\{x_0, y_0, z_0\}$ be the sphere center. For a specific radius R, every point $\{x, y, z\}$ on the sphere satisfies $(x - x_0)^2 + (y - y_0)^2 + (z - z_0)^2 = R^2$. Thus, define

$$c = \begin{bmatrix} x_1^2 + y_1^2 + z_1^2 \\ \vdots \\ x_J^2 + y_J^2 + z_J^2 \end{bmatrix}, A = \begin{bmatrix} 2x_1 & 2y_1 & 2z_1 & 1 \\ \vdots & \vdots & \vdots & \vdots \\ 2x_J & 2y_J & 2z_J & 1 \end{bmatrix}, x = \begin{bmatrix} x_0 \\ y_0 \\ z_0 \\ R^2 - x_0^2 - y_0^2 - z_0^2 \end{bmatrix}$$

$\{x_0, y_0, z_0\}$, R can be found by solving the linear equation $Ax = c$.

By construction, Gag assembly should be part of a sphere with radius 50nm, so R should be very close to 50.

2. Determine the position and size of the ideal spherical cap.

$C_{ideal}$ is on the same sphere as each $p_i$. The surface area of the complex is $S_{complex} = J \times S_{Gag}$, where $S_{Gag}$ is the surface area of one Gag molecule. The surface area of the ideal cap is thus set the same, $S_{ideal} \equiv S_{complex}$. The center of $C_{ideal}$'s base, indicated by $c_C$, should be placed at the COM of the complex, so $c_C = \frac{1}{J}\sum_{i=1}^{J} p_i$. The coverage of $C_{ideal}$ is then only determined by its polar angle $\theta_c = \cos^{-1}(1 - \frac{S_{ideal}}{2\pi R^2})$.

3. Compare the polar angle of each point in P, indicated as $\theta_{p_i}$, with $\theta_c$. RI is the fraction of all monomers within the spherical cap, where H is the Heaviside function and returns 1 only when $\theta_c \geq \theta_{p_i}$:

$$RI = \frac{1}{J} \sum_{i=1}^{J} H(\theta_c - \theta_{p_i}) \qquad (S7)$$

<u>**Supplemental Figures:**</u>

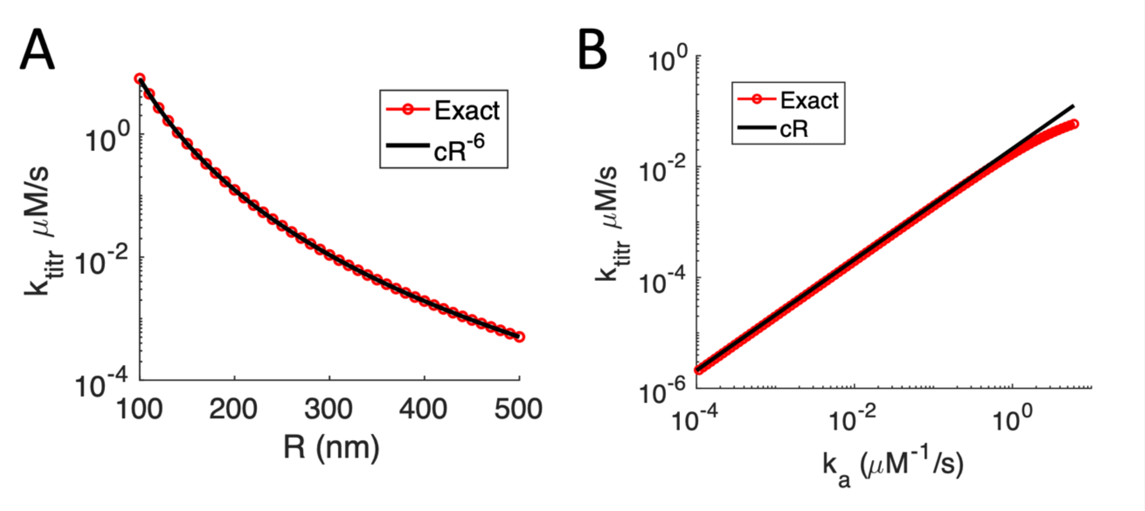


**Figure S1. Derived titration rate is sensitive to changes in volume and binding constants**. A) Our derived titration rate, $k_{\mathrm{titr}}$, Eq. 8 in the main text, scales with $R^{-6}$, as $\tau$ scales with $R^3$ when $R \gg a$. Here we used the model parameters from the main text, and $k_a$ =0.6$\mu$M$^{-1}$s$^{-1}$. B) $k_{\mathrm{titr}}$ increases linearly with $k_a$ over a large range, until binding is limited by diffusion times. Here *R*=300nm. $k_{\mathrm{titr}}$ has an identical dependence on $\rho_0$ as on $k_a$. It also has a power-law dependence on $a$ ($a^{\sim 1.5}$). $k_{\mathrm{titr}}$ has a weaker dependence on $D$, increasing linearly with $D$ only if $Da << k_a$.

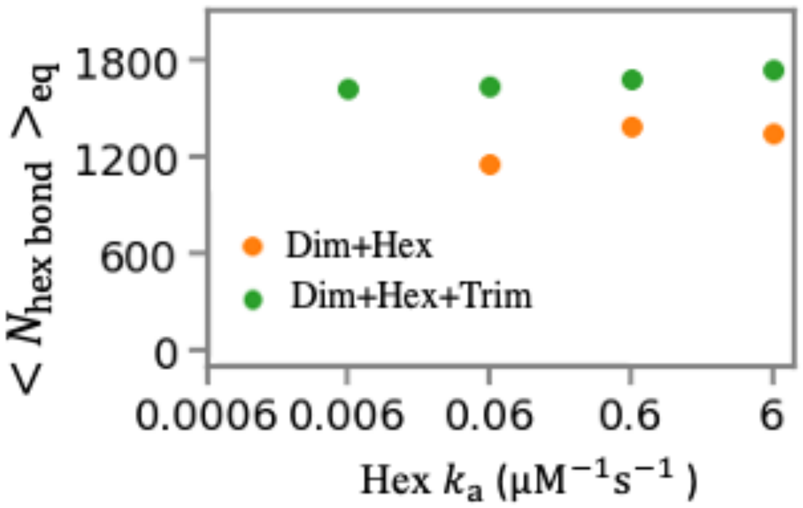


**Figure S2. Validating that the number of equilibrium hexamer bonds is conserved given a fixed free energy, even as rates are increased.** We keep $\Delta G_{hex} = -6.4\ k_BT$ constant but increase the association rate from $k_{a,hex} = 6 \times 10^{-4}$ to $6\ \mu M^{-1}s^{-1}$ and increase dissociation rates accordingly. The number of hexamer bonds formed after steady-state is reached (~100 s) is consistent, although some noise is present due to stochasticity. For the slower rates, the system had not yet reached the equilibrium steady-state. When weak trimer interaction are turned on (green data), $k_{a,trim} = 6.02 \times 10^{-5}\ \mu M^{-1}s^{-1}$, $k_{b,trim} = 1 s^{-1}$, $\Delta G_{trim} = -4.1\ k_BT$, 15% more bonds are formed than without the trimer interaction, $\Delta G_{trim} = 0$ (green data). Dimer interaction are present with $\Delta G_{dim} = -15.6\ k_BT$. All simulations have $D_t = 10\ \mu m^2 s^{-1}, D_{rot} = 0.01\ rad^2 s^{-1}, \Delta t = 0.1\ \mu s$, boxlength=405 nm.

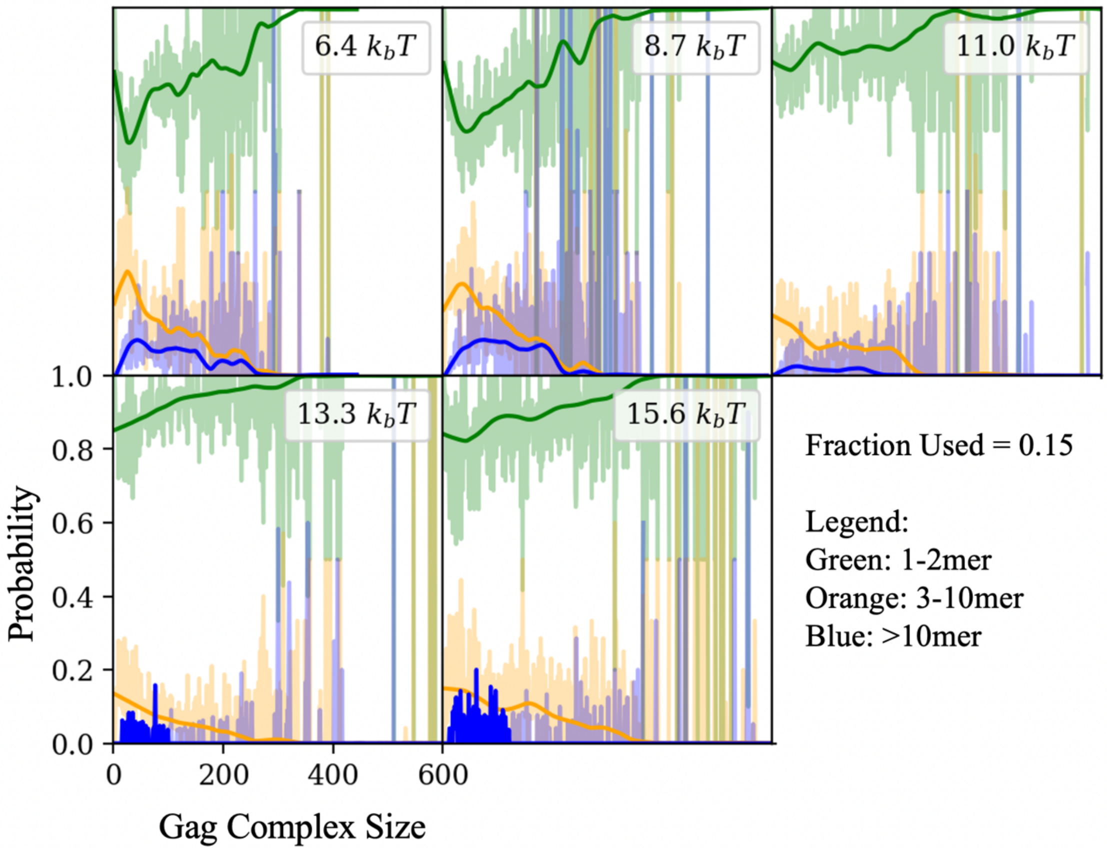


**Figure S3. Growth proceeds largely via addition of monomers and dimers, but larger annealing events also contribute.** Along the x-axis are the sizes of complexes being added, with the y-axis reporting the probability that this complex experiences growth due to addition on monomers/dimers (green), small oligomers (orange) containing less than 10 Gags**,** or larger oligomers (blue)**.** We note that the x-axis indicates the size of the larger complex in the binding instance. For example, if a 20-mer binds to a 5-mer, the graph would show a point at x=20 in the orange category, but this event does not show up for x=5. Thus each binding event will only be counted once. The lines show data smoothing using the LOWESS (Locally Weighted Scatterplot Smoothing) procedure as implemented in the Python statsmodels library, with fraction 0.15. The analysis was performed on simulations with different hexamer contact strength: $\Delta G_{hex} = -6.4, -8.7, -11, -13.3, -15.6\ k_BT$. Annealing between two larger-sized complexes is rare (blue). Larger complexes must have compatible edges and be aligned relatively close to the correct orientation, as significant rotations of large rigid bodies are unphysical over the length of a time-step and are rejected by NERDSS using a threshold (see Methods).

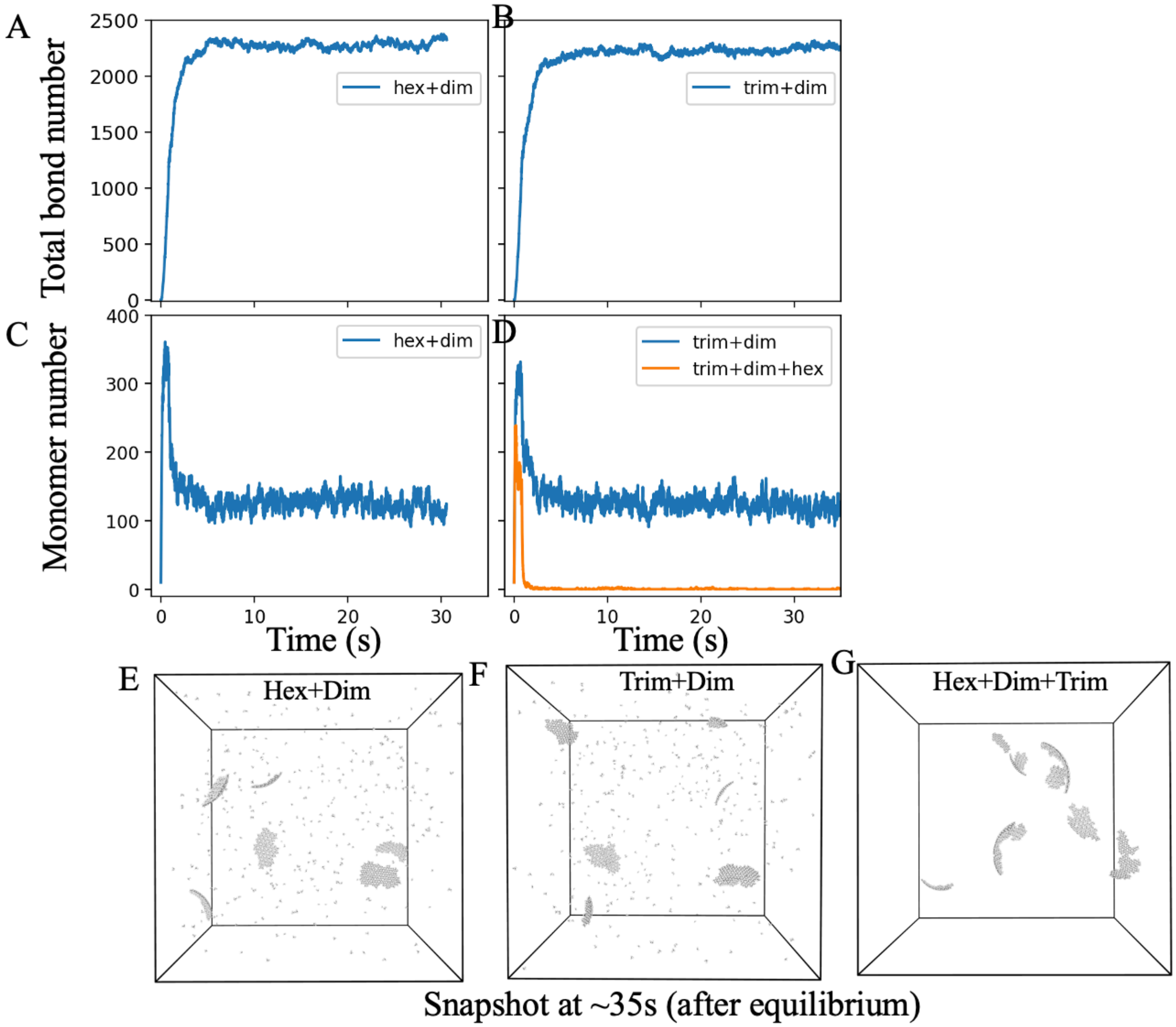


**Figure S4. Phase separation into a dilute phase and incomplete assemblies occurs with only two interactions turned on, conditional on one being relatively weak.** A-B) The total bonds formed in the lattice at equilibrium are similar given the dimer interaction and either the hexamer $\Delta G_{hex} = -6.4\ k_BT$ (A) or trimer (B) $\Delta G_{trim} = -6.4\ k_BT$ contact turned on. $\Delta G_{dim} = -13.3\ k_BT$. C-D), These systems with only two interactions have monomers left over, whereas the addition of the third interaction starves the system of monomers (orange line). E,F,G) are the snapshots at ~35s, where the monomers are visible in (E-F), but fully depleted in (G).

Simulations use $k_{a,dim} = 0.602\ \mu M^{-1}s^{-1}$, $k_{b,dim} = 1s^{-1}$, $\Delta G_{dim} = -13.3\ k_BT$; $k_{a,hex} = k_{a,trim} = 0.602\ \mu M^{-1}s^{-1}$, $k_{b,hex} = k_{b,trim} = 1000s^{-1}$. $\Delta t = 0.5\ \mu s$ and $D_t = 10\ \mu m^2s^{-1}$, $D_{rot} = 0.01\ rad^2s^{-1}$, $\Delta t = 0.5\ \mu s$, water box size 405 nm, $k_c = 60\ \mu M/s$.

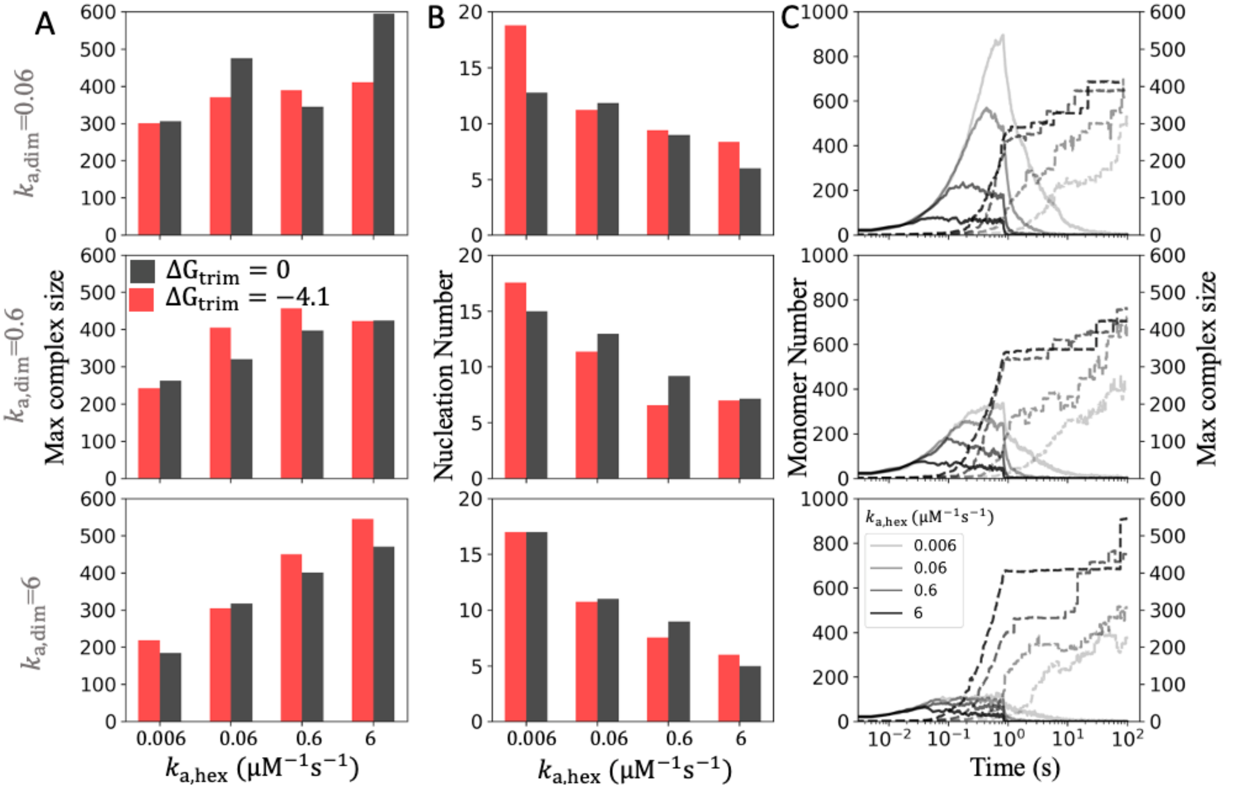


**Figure S5. Faster binding kinetics reduces multiple nucleation events prior to trapping.** A) The size of the largest assembly formed tends to increase as the hexamer association rate increases from 0.006 to 6 $\mu M^{-1}s^{-1}$. From the top row to the bottom row, the dimer association rate $k_{a,dim}$, increases, keeping the off-rate constant. In each subplot, black bars have no trimer interaction, and red bars have $\Delta G_{trim} = -4.1\ k_BT$. B) Correspondingly, the number of nuclei formed decreases as the hexamer rate increases, where a nucleus contains at least 50 monomers. With high $k_{a,hex}$ and $k_{a,dim}$, fewer and larger nuclei are formed. C) The kinetics over 100s shows how monomers are depleted as they are titrated into the system at $k_c = 60\ \mu M/s$, where we stop titration at 0.83s or 50 μM, and trimer interactions are turned on (Left axis, solid curves). We show the growth of the largest complex using dashed lines, on the right axis. These systems all have stable interactions that starve the system of small dimers and oligomers and lead to trapped intermediates, with $\Delta G_{hex} = -8.7, -11.0, 13.3, -15.6\ k_BT$. All simulations included have $k_{b,hex} = 1\ s^{-1}$; $k_{b,dim} = 1\ s^{-1}, \Delta G_{dim} = -11.0, 13.3, -15.6\ k_BT$. $\Delta t = 0.5\ \mu s$ and $D_t = 10\ \mu m^2 s^{-1}, D_{rot} = 0.01\ rad^2 s^{-1}$, boxlength $= 405$ nm.For those with trimer interaction , $k_{a,trim} = 6.02\ \times 10^{-5}\ \mu M^{-1} s^{-1}, k_{b,trim} = 1 s^{-1}$.

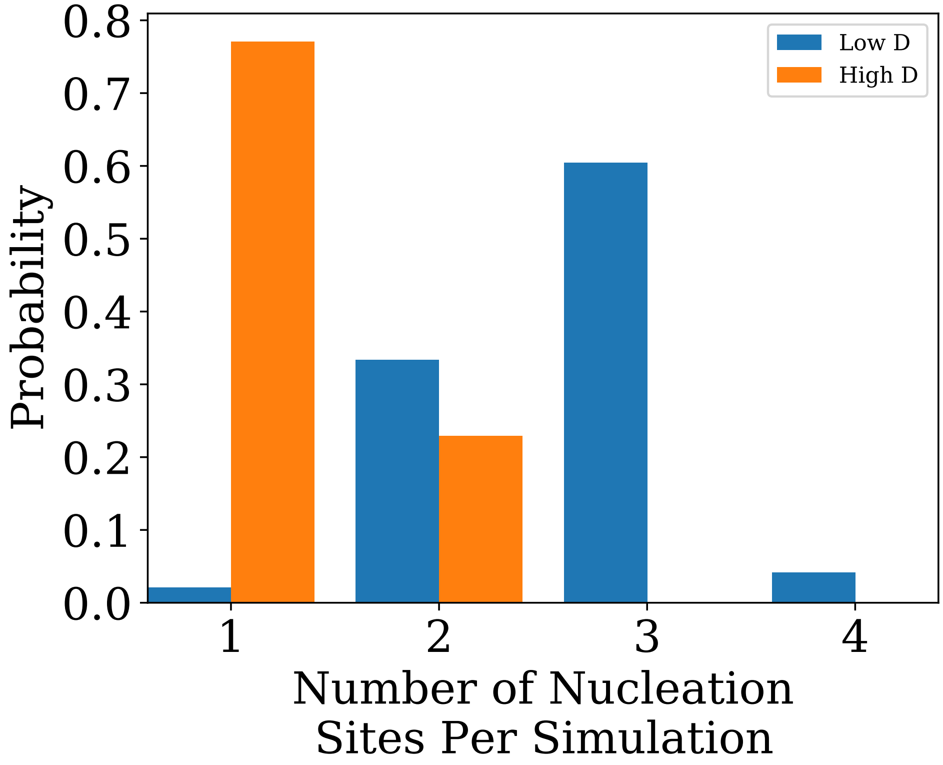


**Figure S6. Higher diffusion coefficients can help to suppress multiple nucleation.** We ran two sets of 48 simulations that differ only in the diffusion coefficient. The plot shows that a faster diffusion (50 vs 10um$^2$/s) promotes slightly fewer nucleations. By testing a faster intrinsic association rate for the slower diffusion simulations and recovering a similar distribution, we expect that the slowed search time is not the primary cause of this result. Instead, NERDSS rejects association events that involve rotations that displace components higher than an average diffusional displacement, times a scale factor. This criteria is thus more permissive to association events involving faster diffusing components. Because these events are only rejected when they involve larger complexes that thus can generate large displacements for monomers distant from the center of rotation, this indicates that there are larger annealing events that help reduce multiple nucleation events forming early on. Overall, this illustrates that larger annealing events can be important in reducing multiple nuclei, and tuning the rarity of these events can thus influence the total number of complexes formed, even though the equilibrium bonds form is preserved.

**SI References:**

[1] Varga, Matthew J. et al. NERDSS: A Nonequilibrium Simulator for Multibody Self-Assembly at the Cellular Scale. Biophysical Journal, Volume 118, Issue 12, 3026 – 3040 (2020).
